# The *Rickettsia* actin-based motility effectors RickA and Sca2 contribute differently to cell-to-cell spread and pathogenicity

**DOI:** 10.1101/2024.08.20.608656

**Authors:** Cuong J. Tran, Zahra Zubair-Nizami, Ingeborg M. Langohr, Matthew D. Welch

## Abstract

*Rickettsia parkeri* is an obligate intracellular, tick-borne bacterial pathogen that can cause eschar-associated rickettsiosis in humans. *R. parkeri* invades host cells, escapes from vacuoles into the cytosol, and undergoes two independent modes of actin-based motility mediated by effectors RickA or Sca2. Actin-based motility of *R. parkeri* enables bacteria to enter protrusions of the host cell plasma membrane that are engulfed by neighboring host cells. However, whether and how RickA and Sca2 independently contribute to cell-to-cell spread *in vitro* or pathogenicity *in vivo* has been unclear. Using live cell imaging of *rickA*::Tn and *sca2*::Tn mutants, we discovered both RickA and Sca2 contribute to different modes of cell-to-cell spread. Compared with Sca2-spread, RickA-spread involves the formation of longer protrusions that exhibit larger fluctuations in length and take a longer time to be engulfed into neighboring cells. We further compared the roles of RickA and Sca2 *in vivo* following intradermal infection of *Ifnar1*^-/-^; *Ifngr1*^-/-^ double-knockout mice, which exhibit eschars and succumb to infection with wild-type *R. parkeri*. We observed that RickA is important for severe eschar formation, whereas Sca2 contributes to larger foci of infection in the skin and dissemination from the skin to the internal organs. Our results suggest that actin-based motility effectors RickA and Sca2 drive two distinct forms of cell-to-cell spread and contribute differently to pathogenicity in the mammalian host.

## Introduction

*Rickettsia* are a genus of Gram-negative obligate intracellular alphaproteobacteria that include multiple groups of pathogenic species (Gillespie et al., 2007; El Karkouri et al., 2022). Spotted fever group (SFG) species represent the largest group and are primarily transmitted by a tick vector. In the Americas, pathogenic SFG species include *R. parkeri* and *R. rickettsii*, the causative agents of milder eschar-associated spotted fever disease (Paddock et al., 2004) and more severe Rocky Mountain spotted fever disease (Dantas-Torres, 2007), respectively. *R. parkeri* in particular has emerged as a model organism to study rickettsial biology under BSL-2 conditions (Scott et al., 2022).

SFG *Rickettsia* are thought to share a common intracellular life cycle. Bacteria invade host cells, escape from a membrane-bound vacuole into the cytosol, polymerize host actin filaments on their surface, and harness the force generated by actin polymerization to drive actin-based motility (Lamason and Welch, 2017). SFG *Rickettsia* genomes encode two effector proteins, RickA and Sca2, that localize to the bacterial surface and mediate actin-based motility (Haglund et al., 2010; Reed et al., 2014). RickA mimics class I nucleation promoting factors (Campellone and Welch, 2010) and directly activates host Arp2/3 complex (Gouin et al., 2004; Jeng et al., 2004), leading to the polymerization of branched actin filament arrays (Gouin et al., 2004; Jeng et al., 2004; Reed et al., 2014). Sca2, in contrast, mimics eukaryotic formin proteins by directly nucleating and elongating unbranched actin filaments (Haglund et al., 2010; Madasu et al., 2013; Alqassim et al., 2019).

Using *rickA*::Tn and *sca2*::Tn transposon mutants to separately define the functions of each protein, it was shown that, in addition to having separate biochemical mechanisms of action, RickA and Sca2 play independent roles in actin-based motility during infection (Reed et al., 2014). *R. parkeri* undergoes two temporally and dynamically different modes of motility during their intracellular lifecycle (Reed et al., 2014; Harris et al., 2018). RickA-driven motility (RickA-motility) occurs early following invasion and escape from the vacuole (<2 h), is slower, and results in curved movement paths. Sca2-driven motility (Sca2-motility) occurs later (>5 h), is faster, and results in straighter trajectories. However, the separate functions of RickA-motility and Sca2-motility during infection *in vitro* and *in vivo* are only beginning to be understood.

One central function of bacterial actin-based motility is to promote cell-to-cell spread through the movement of bacteria to the cell cortex and into protrusions of the plasma membrane that are engulfed into neighboring cells (Lamason and Welch, 2017). Both *rickA*::Tn and *sca2*::Tn mutants have been shown to be attenuated in spread as assessed by plaque size and infectious focus size assays (Kleba et al., 2010; Reed et al., 2014). Furthermore, live cell imaging of cell-to-cell spread of wild-type (WT) *R. parkeri* at one day post infection has shown bacteria undergoing motility and becoming positioned at the plasma membrane, where they enter short protrusions that are engulfed into neighboring cells (Lamason et al., 2016). However, live cell imaging studies have not been reported for *rickA*::Tn or *sca2*::Tn mutants, so it remains unclear how RickA-motility and Sca2-motility independently contribute to cell-to-cell spread.

The contributions of RickA and Sca2 to virulence during SFG *Rickettsia* infection are also poorly understood. A *sca2*::Tn mutant strain of *R. rickettsii* was attenuated following intradermal (i.d.) infection in a guinea pig model (Kleba et al., 2010), and a *sca2*::Tn mutant strain of *R. parkeri* was attenuated following i.d. infection in a *Ifnar1^-/-^; Ifngr1^-/-^* double-knockout (DKO) mouse model (Burke et al., 2021), suggesting that Sca2 is a virulence factor. Studies in the mouse model also suggested that Sca2 is involved in dissemination from the skin into the internal organs (Burke et al., 2021), though the role of Sca2 in tissue-level pathology was not investigated. The role of RickA in animal infection has not yet been reported.

In this study, we used *rickA*::Tn and *sca2*::Tn mutants to assess their individual contribution of both actin-based motility effectors to cell-to-cell spread *in vitro* and pathology *in vivo*. Using live cell imaging, we discovered that RickA-motility propels bacteria into longer and more dynamic protrusions, whereas Sca2-motility positions bacteria at plasma membranes where they then enter into shorter and less dynamic protrusions. Furthermore, using intradermal infections of *Ifnar1*^-/-^; *Ifngr1*^-/-^ DKO mice, we found that Sca2 contributes to colonization of and spread within internal organs, whereas RickA contributes to eschar formation on the skin. Altogether, our results suggest that the two SFG *Rickettsia* actin-based motility effectors, RickA and Sca2, contribute differently to cell-to-cell spread and pathogenicity.

## Results

### The *rickA*::Tn mutant is attenuated in cell-to-cell spread

Previous studies suggested a potential role for RickA-driven motility in cell-to-cell spread, but with inconsistent results (Reed et al., 2014; Lamason et al., 2018). We sought to further clarify the role of *rickA* in spread by assessing the phenotype of the *rickA*::Tn mutant using plaque and infectious focus size assays, which serve as a measure of bacterial replication and cell-to-cell spread. Plaque assays were performed by infecting monolayers of African green monkey kidney epithelial (Vero) cells and observing plaques at 5 d post infection (dpi). We observed *rickA*::Tn bacteria form significantly smaller plaques (1.6x smaller) compared to wild-type (WT) bacteria (Fig. 1A through C). This is consistent with the isolation of *rickA*::Tn bacteria in a screen for mutants that caused a small-plaque phenotype (Lamason et al., 2018). Infectious focus size assays were performed by infecting monolayers of human lung epithelial (A549) cells and visualizing foci at 2 dpi. We observed *rickA*::Tn bacteria form significantly smaller foci (1.4x fewer cells per focus) than WT bacteria (Fig. 1D through F), consistent with earlier results (Reed et al., 2014). The observed differences in plaque and infectious focus sizes between *rickA*::Tn and WT bacteria were not due to differences in growth rates, as it was previously shown that *rickA*::Tn and WT bacteria grew at similar rates in Vero cells (Reed et al., 2014). Altogether, these data suggest that RickA-driven motility is important for cell-to-cell spread.

**Figure 1.**
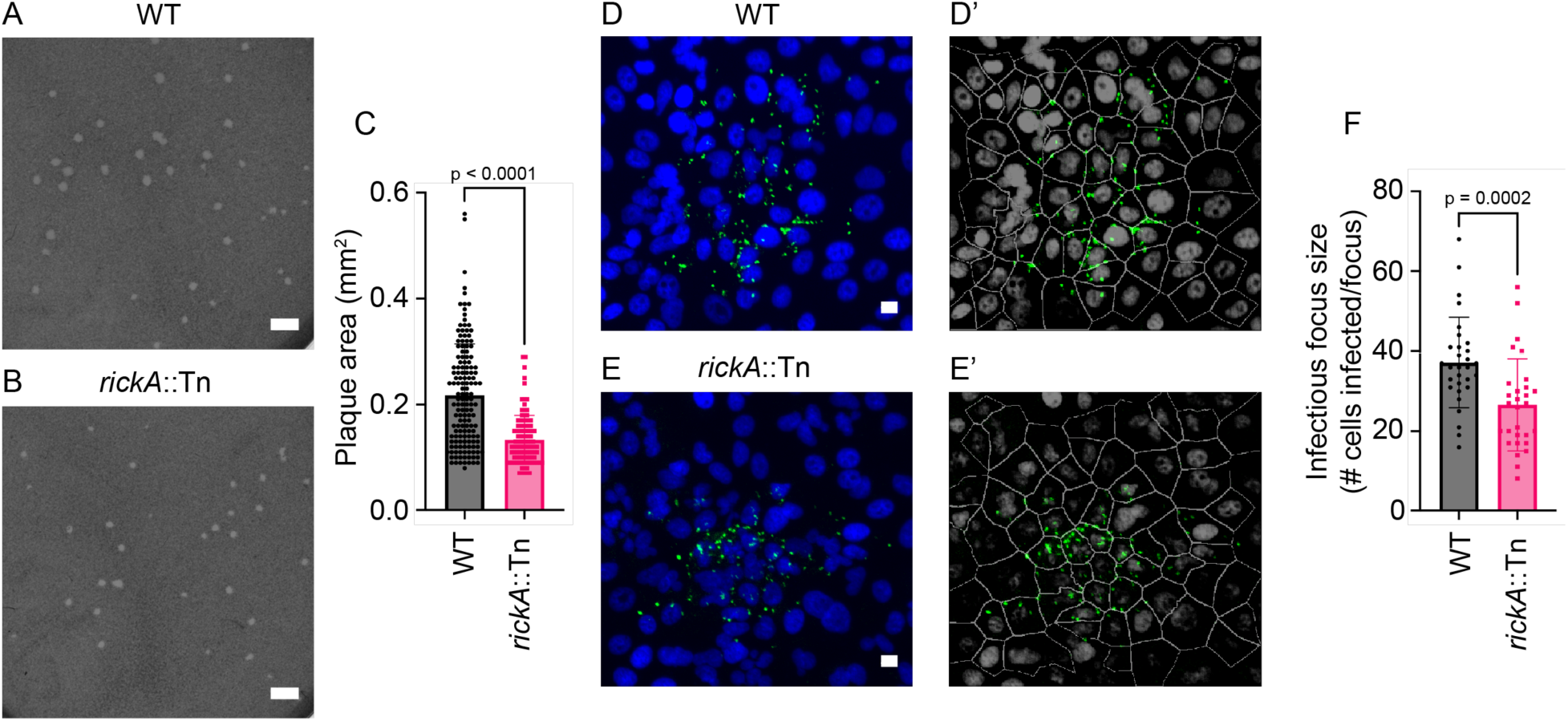
*rickA*::Tn bacteria form smaller plaques and infect fewer cells per infectious focus compared with WT bacteria. **(A-B)** Representative image of plaques in Vero cell monolayers at 5 dpi with (A) WT or (B) *rickA*::Tn bacteria. **(C)** Quantification of plaque diameters from A and B (n=2 biological replicates with 45-76 plaques quantified per replicate). **(D-E)** Representative images of infectious foci in monolayers of A549 cells at 2 dpi with (D) WT bacteria at an MOI of 0.005 or (E) *rickA*::Tn bacteria at an MOI of 0.02. Nuclei were stained with DAPI (blue) and bacteria were stained with α-*Rickettsia* 14-13 (green). **(D’-E’)** Images from CellProfiler analysis showing nuclei and cell borders (grey), as well as bacteria (green). **(F)** Quantification of number of infected A549 cells per infectious focus in A and B (n=3 independent experiments per strain with 10 infectious foci analyzed per experiment). Scale bars in (A) and (B) are 2 mm; scale bars in (D) and (E) are 10 μm. Error bars are mean ± SD. p-values in C and F are from two-tailed Mann-Whitney tests.

### Both RickA-motility and Sca2-motility directly contribute to cell-to-cell spread

A previous study documenting the role of actin-based motility in cell-to-cell spread of *R. parkeri* examined this process for WT bacteria only and thus was unable to parse any separate roles for RickA-motility versus Sca2-motility in spread (Lamason et al., 2016). To determine the separate roles of RickA versus Sca2 in spread, we made use of *sca2*::Tn bacteria, which are only able to undergo RickA-motility, and *rickA*::Tn bacteria, which are only able to undergo Sca2-motility. Mutant bacteria expressing GFPuv from the transposon insertion were infected into human lung epithelial A549 cells expressing a plasma membrane marker, TagRFP-t-farnesyl, as well as the filamentous-actin (F-actin) marker, LifeAct-3xTagBFP (Lamason et al., 2016). This approach allowed for the simultaneous visualization of bacteria, F-actin, and host cell plasma membranes in live cell imaging experiments.

We first tested whether RickA-motility directly contributes to spread by imaging cells infected with *sca2*::Tn mutant commencing at ∼5-10 min post infection (mpi), when RickA-motility begins (Reed et al., 2014). We observed that bacteria undergoing RickA-motility were propelled against the plasma membrane of the infected “donor” cell and pushed into long protrusions of the plasma membrane that extended into neighboring “recipient” cells (Fig. 2). Bacteria within protrusions maintained their actin tails. In 12 of 26 examples, protrusions containing a bacterium were engulfed into recipient cells and resolved into secondary vacuoles, resulting in successful spread to the neighboring recipient cell (Movie S1). In 14 of 26 examples, protrusions containing bacteria retracted back into the donor cell (Movie S2), resulting in a failure to spread to the neighboring cell. Overall, these observations demonstrate a role for RickA-driven motility in mediating cell-to-cell spread.

**Figure 2.**
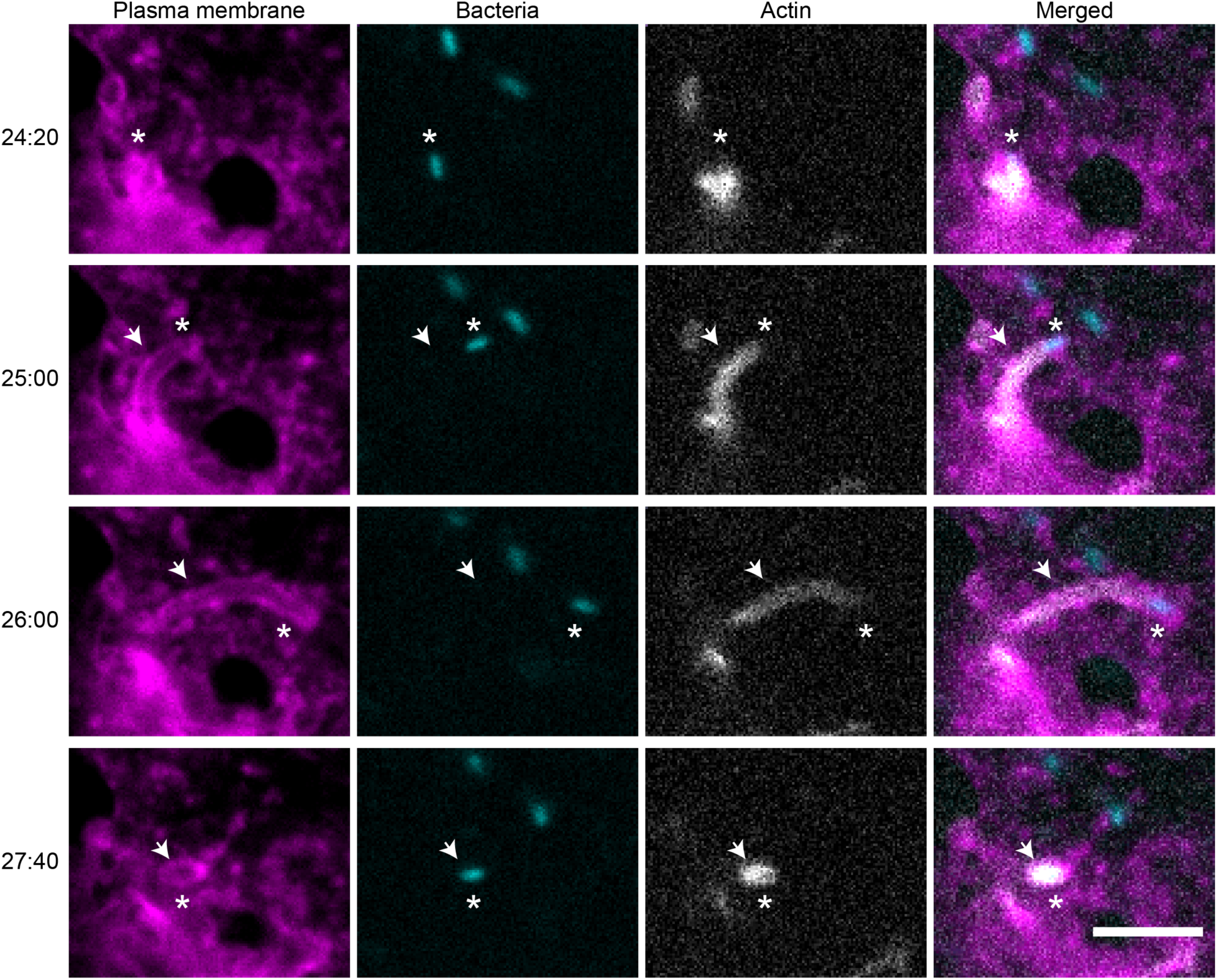
RickA-motility mediates cell-to-cell spread. Still images from a video of A549 cells expressing TagRFP-t-farnesyl (magenta) and LifeAct-3xTagBFP (grey) infected with *sca2*::Tn bacteria (cyan) starting at 5-10 mpi. Timestamps on the left are MM:SS. Asterisks indicate location of a bacterium undergoing actin-based motility and cell-to-cell spread. Arrows indicate the location of a protrusion. Scale bar is 5 μm.

We next tested the role of Sca2-motility in spread by imaging cells infected with the *rickA*::Tn mutant starting at 27 h post infection (hpi) when Sca2-motility is prevalent (Reed et al., 2014) and cell-to-cell spread of WT bacteria had previously been seen (Lamason et al., 2016). We observed bacteria undergoing Sca2-driven motility being pushed to the plasma membrane and then becoming immotile for a brief period before entering into short protrusions of the donor cell plasma membrane (Fig. 3, Movie S3). In all 10 of 10 examples, protrusions were engulfed into neighboring recipient cells (though the short protrusions generated during Sca2-spread were harder to definitively image, making it possible that failed cell-to-cell events were not captured). Additionally, we imaged A549 cells infected with *sca2*::Tn bacteria at 27 hpi but failed to observe cell-to-cell spread events driven by RickA-motility at this timepoint. Taken together, these results demonstrate that RickA-motility contributes to spread primarily at earlier times, and Sca2-motility contributes to spread at later times during infection.

**Figure 3.**
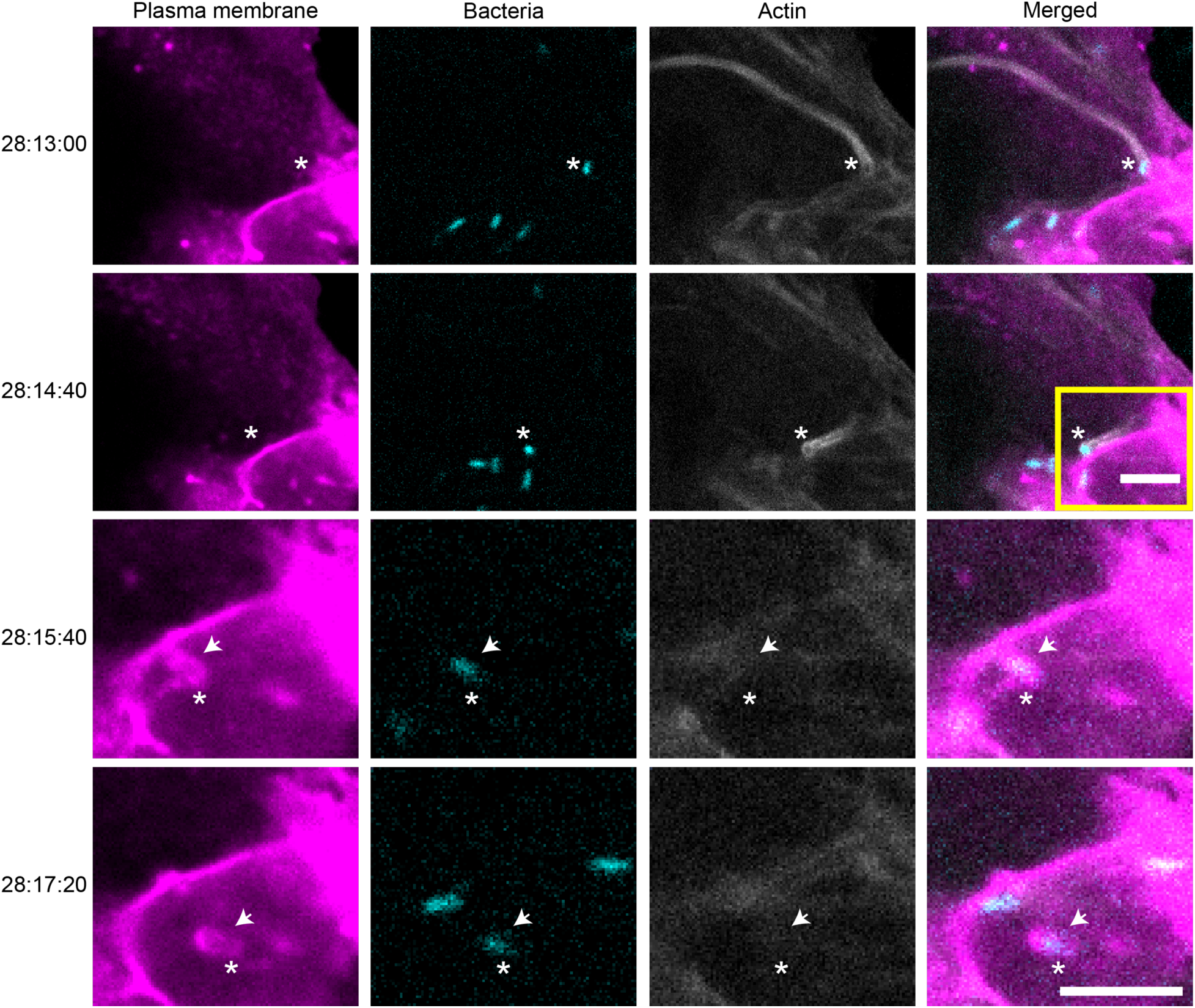
Sca2-motility mediates cell-to-cell spread. Still images from a video taken of A549 cells expressing TagRFP-t-farnesyl (magenta) and LifeAct-3xTagBFP (grey) infected with *rickA*::Tn bacteria (cyan) starting at 28 hpi. Timestamps on the left are HH:MM:SS. Asterisks indicate location of a bacterium undergoing actin-based motility and cell-to-cell spread. Arrows indicates the locations of a protrusion. The yellow box indicates a region that was magnified for the bottom rows with timestamps 28:15:40 and 28:17:20. Scale bars for top two rows and bottom two rows are both 5 μm.

### The dynamics of RickA-driven and Sca2-driven cell-to-cell spread differ

We next sought to further describe and compare the dynamics of RickA-driven versus Sca2-driven cell-to-cell spread. We quantified parameters including time spent within protrusions, protrusion lengths, and bacterial movement within protrusions. To compare time spent within protrusions for RickA-spread versus Sca2-spread, we measured the time elapsed between protrusion initiation (defined as the moment a bacterium entered an indentation in the donor cell plasma membrane) and protrusion engulfment (defined as the moment a protrusion resolved into secondary vacuole in the recipient cell). We found that the time spent in protrusions was significantly longer (by 2.6x) for RickA-spread versus Sca2-spread (Fig. 4A).

**Figure 4.**
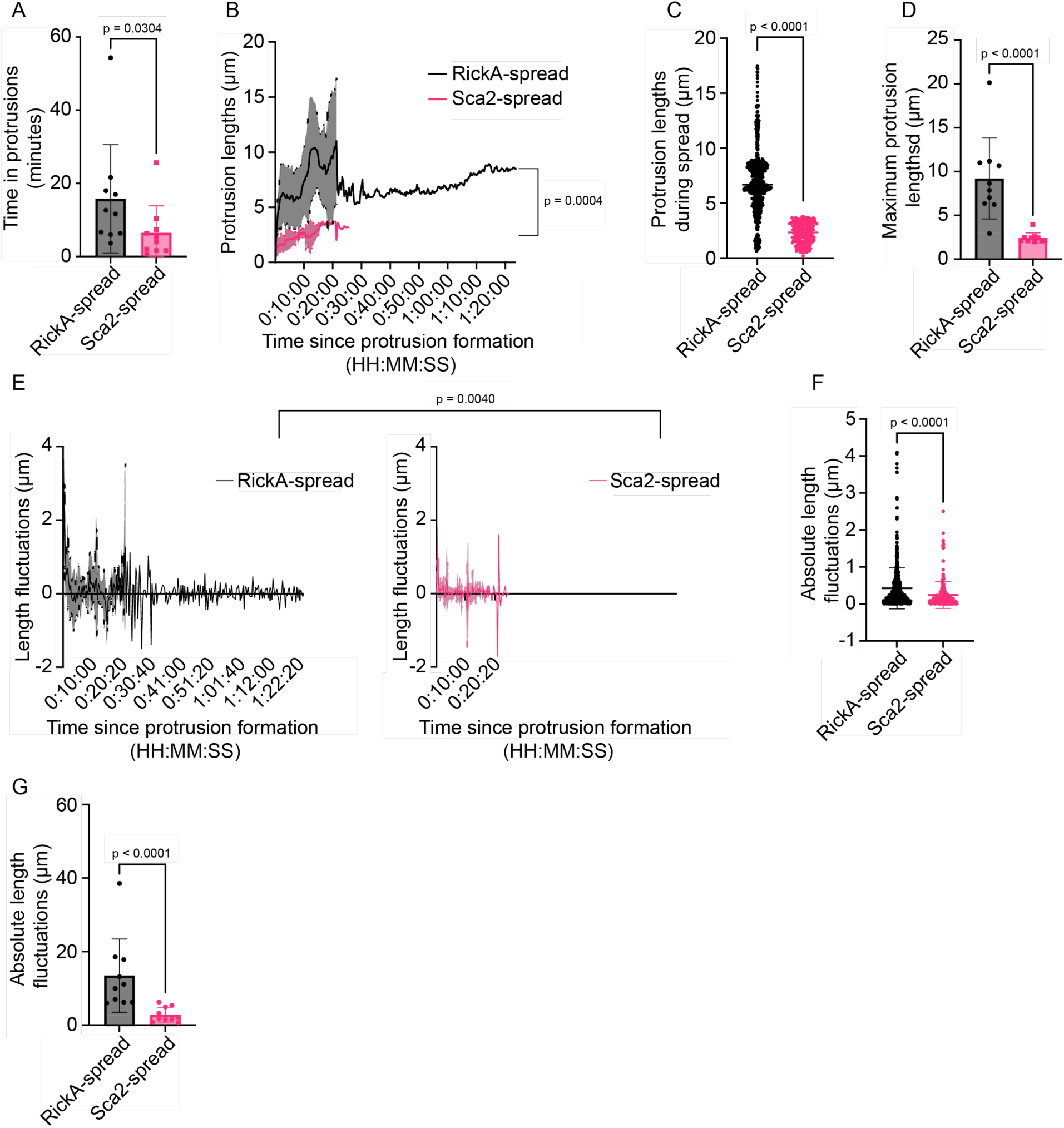
RickA-driven and Sca2-driven cell-to-cell spread exhibit different dynamics. **(A)** Time spent within protrusions for RickA-spread (black) versus Sca2-spread (pink). **(B)** Average protrusion length over time during RickA-spread (black) versus Sca2-spread (pink). **(C)** All protrusion lengths during RickA-spread (black) and Sca2-spread (pink). **(D)** Maximum protrusion lengths during RickA-spread (black) versus Sca2-spread (pink). **(E)** Comparison of the fluctuation in the length of protrusions during RickA-spread (black) versus Sca2-spread (pink) taken at consecutive 20 s intervals. **(F)** Quantification of the absolute value of the length fluctuations between each consecutive position at 20 s intervals during RickA-spread (black) versus Sca2-spread (pink). **(G)** Total distance bacteria traveled during RickA-spread (black) versus Sca2-spread (pink). (A-F) n = 10 cell-to-cell spread events from 8 – 10 independent experiments. Data (A, C, D, and G) are mean ± SD. Thick lines in (B) and (E) represent mean values while shaded regions represent mean ± SD. p-values for (A), (C), (D) and (F-G) are from two-tailed Mann-Whitney test. p-values in (B) and (E) are from a mixed-effects analysis.

We also measured the length of protrusions over time. In general, protrusions elongated to an initial length, then underwent cycles of retraction and further elongation, fluctuating in length over time (Movie S4). We measured the straight-line distance between the protrusion origin and the protrusion tip, at 20 s intervals, over the entire time course from protrusion initiation to engulfment, for 10 examples each of RickA-driven versus Sca2-driven spread (Fig. 4B, Fig. S1). The average protrusion lengths over time were significantly different for RickA-spread versus Sca2-spread (Fig. 4B and C). We also measured the maximum length of protrusions attained during the time interval by measuring the arc lengths (distance between two points along a curve) from the proximal base to the distal tip of each protrusion. The maximum lengths of protrusions generated during RickA-spread were significantly longer (by 4x) than protrusions generated during Sca2-spread (Fig. 4D).

Finally, we compared the dynamic behavior of RickA-generated and Sca2-generated protrusions by assessing the fluctuation in the position of the tips of protrusions during spread. We quantified differences in the straight-line protrusion lengths between consecutive 20 s intervals over the course of spread, with positive numbers indicating elongation, and negative numbers indicating retraction (Fig. 4E). We observed a significant difference, with RickA-generated protrusions exhibiting larger fluctuations in length than Sca2-generated protrusions (Fig. 4F). We also quantified the total distance traveled by individual bacteria as they fluctuated in position during RickA-spread and Sca2-spread by summing the absolute value of the changes in position of the protrusion tip between each 20 s interval. Our data revealed that bacteria traveled an average total distance of 34 ± 25 μm during RickA-spread, whereas bacteria traveled an average distance of 7 ± 5 μm during Sca2-spread (Fig. 4G). Altogether, these data show that the parameters and dynamics between RickA-spread and Sca2-spread are different.

### The *rickA*::Tn mutant is not attenuated in lethal infection *in vivo*

The observation that RickA and Sca2 contribute to different modes of cell-to-cell spread, with different timing and parameters, raised the question of how each contributes to infection *in vivo*. We previously reported that *Ifnar1^-/-^; Ifngr1^-/-^* DKO mice, carrying mutations in the genes encoding the receptors for IFN-I (*Ifnar1*) and IFN-γ (*Ifngr1*), are susceptible to eschar-associated rickettsiosis (Burke et al., 2021). Using this animal model, we previously showed that, following i.d. infection, *sca2*::Tn bacteria are attenuated in lethality (Burke et al., 2021).

To test the importance of RickA in pathogenesis, we infected *Ifnar1^-/-^; Ifngr1^-/-^* DKO mice with 1×10^3^ pfu of either WT or *rickA*::Tn mutant *R. parkeri*. During a 35-day time course, mice infected with WT or *rickA*::Tn strains had similar survival rates (Fig. 5A), body temperature profiles (Fig. 5B), and changes in body weight (Fig. 5C). Thus, we did not detect a role for RickA in lethal infection.

**Figure 5.**
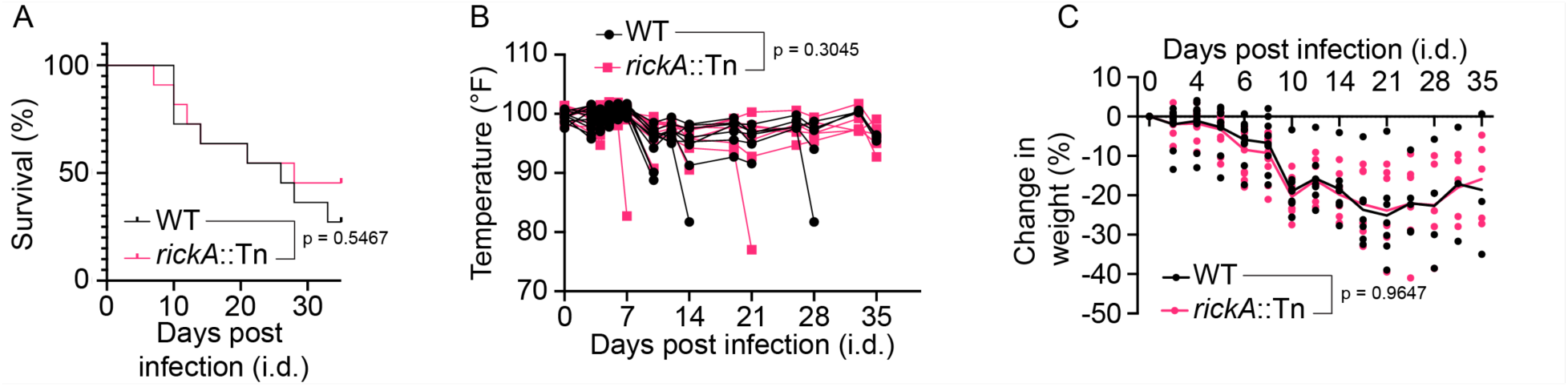
RickA is not required for lethal infection. **(A)** Survival curve following i.d. infection of mice with WT (black) versus *rickA*::Tn bacteria (pink) over a 35 d time-course. **(B)** Temperature changes over time following i.d. infection of mice with WT (black) versus *rickA*::Tn bacteria (pink) over a 35d time-course. Each line represents an individual mouse. **(C)** Changes in percent weight following i.d. infection of mice with WT (black) versus rickA::Tn (pink) bacteria. All data came from n = 11 mice per strain from 3 independent experiments. Each dot represents a single mouse. Solid line represents average change in weight. p-value in (A) was from Mantel-Cox test; p-values for (B) and (C) are from mixed-effects analyses.

### The *sca2*::Tn mutant, but not the *rickA*::Tn mutant, is significantly attenuated in colonization of internal organs

We previously showed that, following i.d. infection of the skin, *sca2*::Tn bacteria are defective in colonizing the liver and spleen compared to WT bacteria (Burke et al., 2021). However, the phenotype of the *rickA*::Tn mutant in dissemination to internal organs was not previously examined. We therefore sought to directly compare the ability of the WT, *sca2*::Tn, and *rickA*::Tn strains to colonize internal organs by assessing the bacterial burden in the skin, spleen, liver, lung, and brain following i.d. infection of *Ifnar1^-/-^; Ifngr1^-/-^* DKO mice. We chose 10 d post infection (dpi) as our endpoint because mice began to succumb to infection at this time. We found that similar numbers of WT, *rickA*::Tn, and *sca2*::Tn bacteria were present in the skin at the site of infection (Fig. 6A). However, *sca2*::Tn mutant bacteria were present in significantly reduced numbers in spleen, liver, lung, and brain tissue, corroborating prior data suggesting that Sca2 is important for colonization of liver and spleen (Burke et al., 2021). In contrast, the numbers of *rickA*::Tn mutant bacteria in internal organs were indistinguishable from WT (Fig. 6B through E). These results confirm a role for Sca2 in dissemination to and colonization of internal organs, but do not reveal a significant role for RickA in this process.

**Figure 6.**
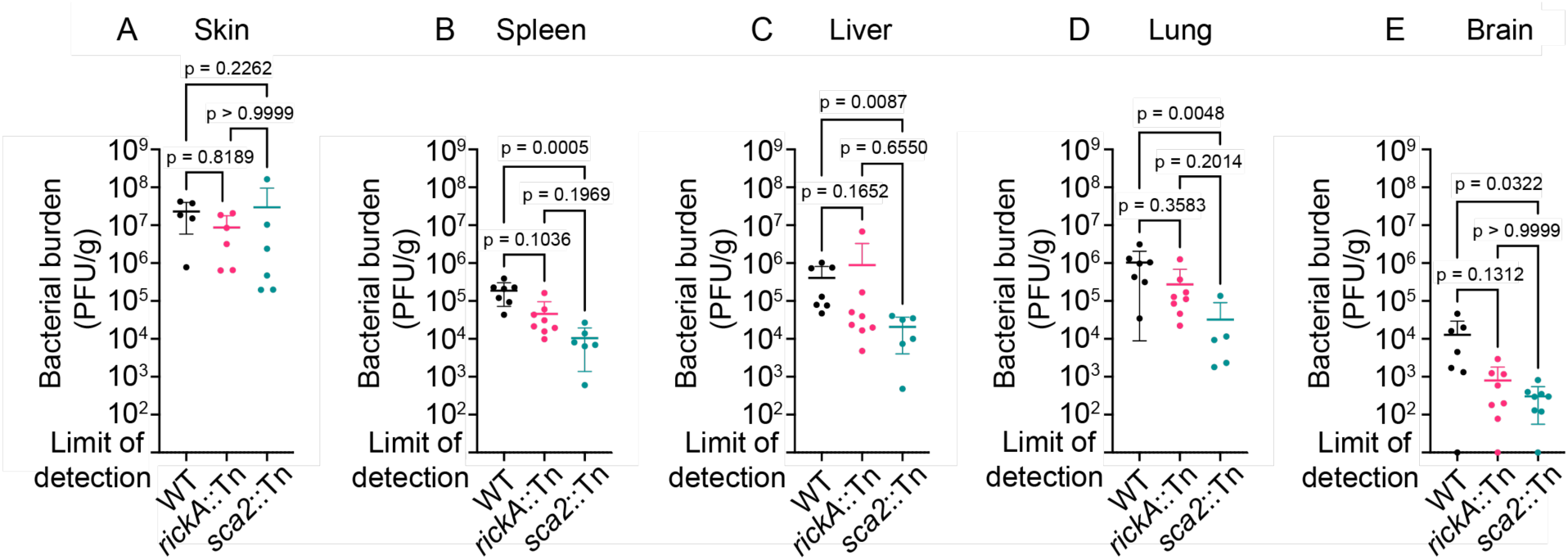
Sca2 contributes to internal organ colonization. Quantification of bacterial burden in the **(A)** skin, **(B)** spleen, **(C)** liver, **(D)** lung, and **(E)** brain of mice following i.d. infection with 1×10^3^ pfu WT (black), *rickA*::Tn (pink) or *sca2*::Tn (blue) bacteria for 10 d. Error bars are mean ± SD. For the skin: n = 5 mice infected with WT, n = 6 for each *rickA*::Tn and *sca2*::Tn strains, data are from 2 independent biological replicates. For the spleen and liver: n = 7 mice for WT, n = 8 for *rickA*::Tn, and n = 6 for *sca2*::Tn, data are from 3 independent experiments. For the lung: n = 7 mice for WT, n = 8 for *rickA*::Tn, and n = 5 for *sca2*::Tn, data are from 3 independent experiments. For brain: n = 7 mice for WT, and n = 8 for *rickA*::Tn- and *sca2*::Tn; data are from 3 independent experiments. p-values are from Dunn’s multiple comparison test.

### Tissue-level pathology was similar for *sca2*::Tn, *rickA*::Tn, and wild-type strains, but *sca2*::Tn showed significantly reduced areas of infection

To determine whether RickA or Sca2 contribute to pathological changes at the tissue level during infection, we performed hematoxylin and eosin (H&E) staining as well as immunohistochemistry (IHC) on skin, spleen, lung, liver, and brain tissue sections from i.d.- infected *Ifnar1^-/-^; Ifngr1^-/-^* DKO mice at 10 dpi with WT, *rickA*::Tn, and *sca2*::Tn strains. Mice infected intradermally with any of the aforementioned strains exhibited similar overall alterations, as well as similar extent and degree of inflammation in all tissues (Fig. 7A through E, Fig. S2 and 3). In the skin, the inflammation was generally most severe in the subcutis, consisting primarily of necrosuppurative exudate, whereas mixed neutrophilic and histiocytic infiltrate variably expanded the deep dermis. Occasional vasculitis and thrombosis were present. Rarely, there was also coagulative (ischemic) necrosis of the dermis and epidermis. In the spleen, histologic alterations consisted primarily of increased extramedullary hematopoiesis in the red pulp and of plasmacytosis in the white pulp, indicative of a systemic cellular and humoral response, respectively, to the rickettsial infection. Rarely, thrombosis and sparse foci of neutrophilic and histiocytic infiltrate were also present. The liver had multifocal to coalescing mixed neutrophilic and histiocytic inflammation, with variable central fibrin exudation, necrosis, and thrombosis. In the lung, there was mixed neutrophilic and mononuclear inflammatory infiltrate, often centered on vessels. The affected vasculature was lined by hypertrophied endothelium, with variable adhered and subendothelial infiltrating leukocytes, accompanied by multifocal vascular thrombosis as well as hemorrhage and edema of the surrounding pulmonary parenchyma. Lastly, in the brain, several of the mice had multifocal, often minimal leptomeningeal and rarely subependymal inflammatory infiltrates, infrequently with thrombosis and fibrin exudation.

**Figure 7.**
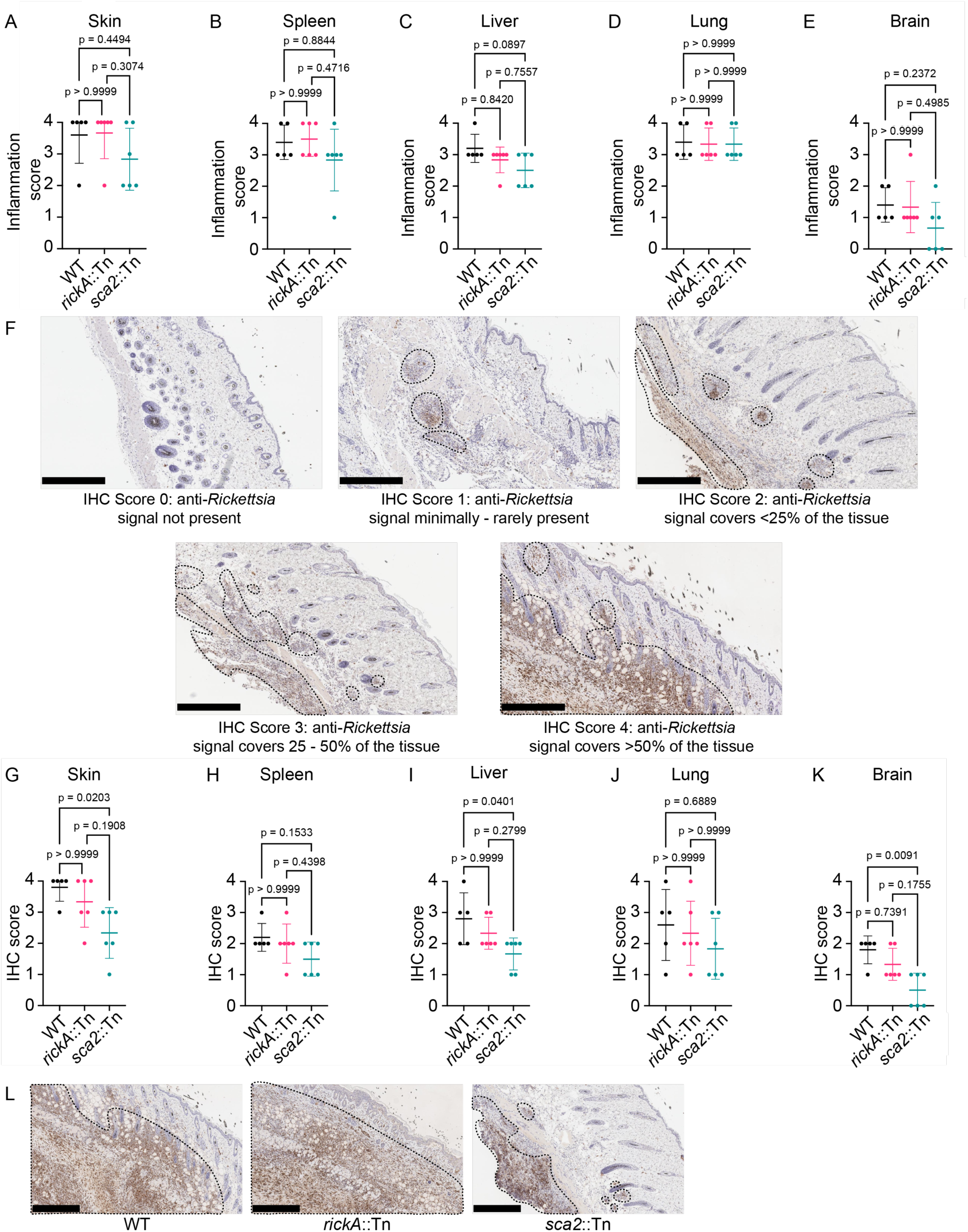
Sca2 contributes to spread in the skin, liver, and brain. Inflammation scores from **(A)** skin, **(B)** spleen, **(C**) liver, **(D)** lung, and **(E)** brain sections of mice infected intradermally with 1×10^3^ pfu WT (black), *rickA*::Tn (pink), or *sca2*::Tn (blue) bacteria for 10 d. **(F)** Representative images for IHC scores. Dotted black lines indicate areas of anti-*Rickettsia* staining (brown). IHC scores from **(G)** skin, **(H)** spleen, **(I)** liver, **(J)** lung, and **(K)** brain sections of mice infected intradermally with 1×10^3^ pfu WT (black), *rickA*::Tn (pink), or *sca2*::Tn (blue) bacteria for 10 d. **(L)** Representative IHC images from the skin of mice infected with WT, *rickA*::Tn, or *sca2*::Tn bacteria for 10 d. Dotted lines areas of anti-*Rickettsia* staining (brown). Error bars are mean ± SD. n = 5 mice infected with WT, n = 6 for *rickA*::Tn and *sca2*::Tn strains, data are from 2 independent experiments. p-values are from Dunn’s multiple comparison test. Scale bars in (F) and (L) are 500 μm.

IHC highlighted the presence of bacteria within inflammatory foci in all mice. Bacteria were present primarily in neutrophils and histiocytes, including in tissue-resident macrophages, as well as in scattered endothelial cells of intralesional vessels. Significantly fewer bacteria were present in the skin, liver, and brain of the *sca2*::Tn infected mice. To quantify the extent of infection in the skin, spleen, liver, lung, and brain, we assigned an IHC score based on the percent area of a given tissue section that was stained with anti-*Rickettsia* antibody (Fig. 7F). We found that *sca2*::Tn bacteria had significantly lower immunohistochemistry scores in the skin (Fig. 7G), liver (Fig. 7I), and brain (Fig. 7K). In contrast, no differences in immunohistochemistry scores between *rickA*::Tn and WT bacteria were observed in any of the tissues examined (Fig. 7G through K). Altogether, these results suggest that Sca2, but not RickA, is important for infecting more extensive tissue areas within the skin and internal organs.

### The *rickA*::Tn mutant causes attenuation of eschar formation in the skin

A key outcome of i.d. infection of *Ifnar1^-/-^; Ifngr1^-/-^* DKO mice with *R. parkeri* is the development of eschars starting at 6 dpi, similar to eschars formed in human patients (Burke et al., 2021). Although i.d. infection with *sca2*::Tn or WT bacteria were previously reported to result in eschars of similar severity, the contribution of RickA to eschar formation was not reported. To assess the role of RickA to eschar formation, we infected mice i.d. with 1×10^3^ WT or *rickA*::Tn bacteria and quantified the severity of eschars over the 35 d time course using a previously developed scoring guide (Burke et al., 2021). Mice infected with *rickA*::Tn bacteria presented with significantly lower eschar scores than mice infected with WT bacteria (Fig. 8A and B), indicating significantly reduced inflammation and scabbing of the skin. Thus, *rickA* is important for eschar formation in the skin.

**Figure 8.**
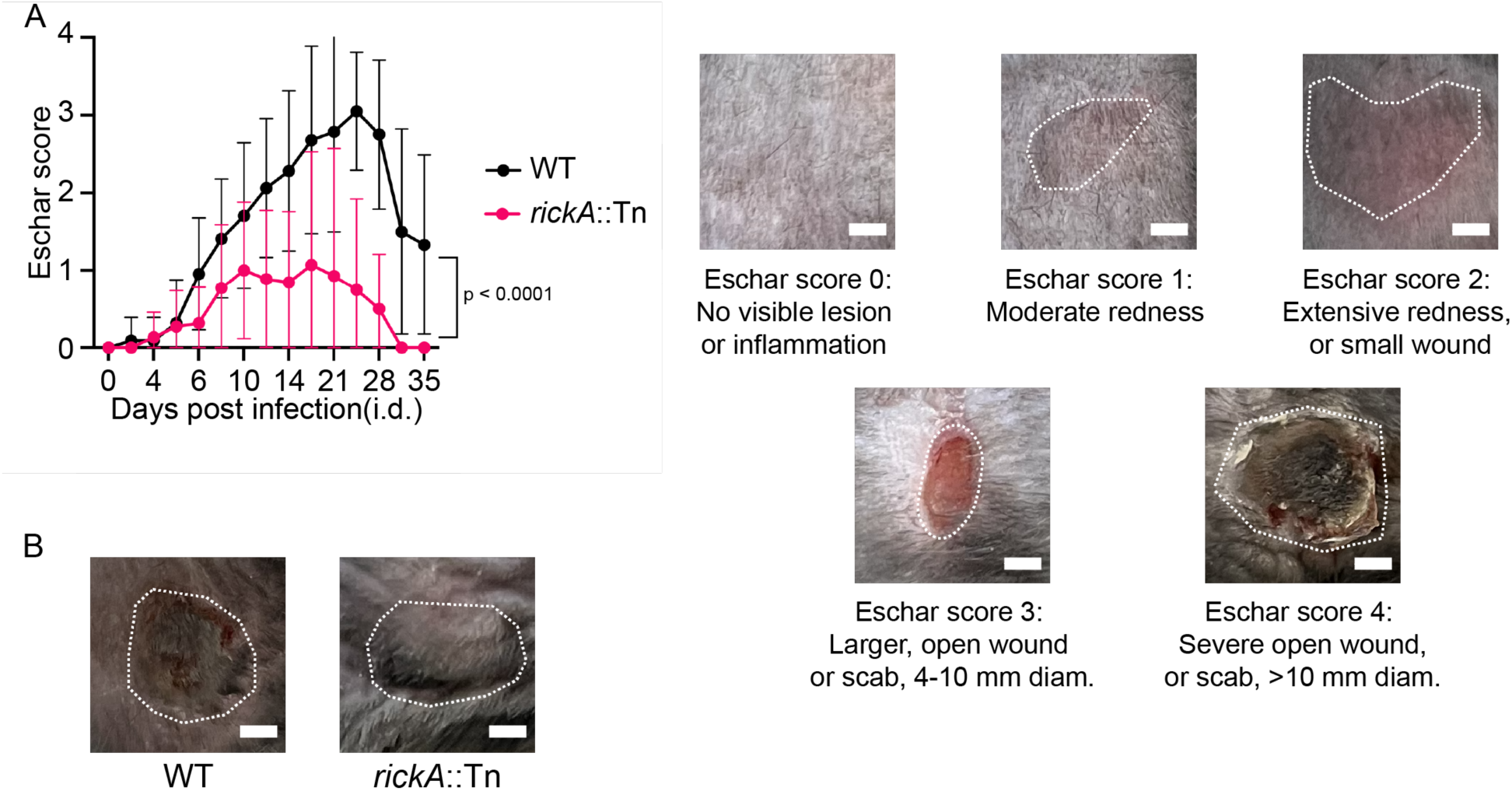
RickA contributes to eschar formation. **(A)** (Left) Changes in eschar scores from mice infected intradermally with WT (black) versus *rickA*::Tn (pink) bacteria over a 35 d time-course. Solid lines represent average eschar scores, and error bars represent mean ± SD. (Right) Representative images of each eschar score. Dotted white lines indicate area of the eschar. **(B)** Representative images of an eschar from a mouse infected with WT bacteria (left) and a mouse infected with *rickA*::Tn bacteria at 19 dpi. Dotted white lines indicate area of the eschar. All data came from n = 11 mice per strain from 3 independent experiments. Scale bars represent 3 mm. p-value is from mixed-effects analysis.

## Discussion

The roles of the two rickettsial actin-based motility effectors, RickA and Sca2, in infection *in vitro* and *in vivo* have remained an enigma. In this study, we demonstrate that both RickA-motility and Sca2-motility play independent roles in mediating cell-to-cell spread. RickA-motility propels bacteria against the host cell plasma membrane to generate longer and more dynamic protrusions, whereas Sca2-motility positions bacteria near the plasma membrane where they enter into shorter and less dynamic protrusions. In a mouse model of infection, RickA is important for severe eschar formation. Sca2, in contrast, is required for lethality, internal organ colonization, and spread within the skin. Altogether, these results suggest that the two modes of SFG *Rickettsia* actin-based motility contribute differently to cell-to-cell spread *in vitro* and pathogenesis *in vivo*.

Whether RickA contributes to cell-to-cell spread had been unclear. In one study, *rickA*::Tn mutants were found to infect fewer cells per infectious focus than WT, but formed plaques of similar size (Reed et al., 2014). In another study, *rickA*::Tn mutants were isolated in a qualitative visual screen for small plaque mutants (Lamason et al., 2018). We confirmed that *rickA*::Tn bacteria infect fewer cells per infectious focus and form smaller plaques than WT *R. parkeri*. These findings are consistent with observations of *Listeria monocytogenes* mutants in the actin-based motility gene, *actA* (Domann et al., 1992; Kocks et al., 1992); and *Shigella flexneri* mutants in the actin-based motility gene, *icsA* (Bernardini et al., 1989), which were unable to form plaques. The observation that *R. parkeri rickA*::Tn or *sca2*::Tn mutants still form plaques (Kleba et al., 2010; Reed et al., 2014; Lamason et al., 2018) suggests that the two proteins may have redundant roles in cell-to-cell spread and plaque formation.

Cell-to-cell spread of WT *R. parkeri* was previously observed to involve actin-based motility that positions bacteria near the plasma membrane of an infected donor cell, where bacteria cease motility and enter into short protrusions that are engulfed into a neighboring recipient cell (Lamason et al., 2016). However, it was unclear whether the observed motility and cell-to-cell spread of WT bacteria was mediated by RickA or Sca2. Using live cell imaging of the *sca2*::Tn mutant, which only undergoes RickA-motility; and the *rickA*::Tn mutant, which only undergoes Sca2-motility, we parsed the contributions of RickA-motility and Sca2-motility to spread. In contrast to what was described for WT *R. parkeri* (Lamason et al., 2016), we observed bacteria at 10-15 mpi undergoing RickA-motility pushing on the donor cell plasma membrane and entering into longer protrusions that exhibited greater amplitudes of oscillations between protrusion and retraction. This form of spread is similar to the “fitful” movements observed from protrusions generated during *L. monocytogenes* cell-to-cell spread (Robbins et al., 1999). The duration of spread was also longer than previously described for WT *R. parkeri* (Lamason et al., 2016). RickA-spread is similar to spread during infection with *L. monocytogenes* (Sechi et al., 1997; Gouin et al., 1999; Robbins et al., 1999; Lamason et al., 2016) and *S. flexneri* (Kadurugamuwa et al., 1991; Gouin et al., 1999; Dragoi and Agaisse, 2014), which involve both long protrusions and durations of spread events compared to what has previously been observed for WT *R. parkeri* infection. Long protrusions are also observed during *Mycobacterium marinum* spread (Stamm et al., 2003). RickA-motility involves activation of the host Arp2/3 complex (Gouin et al., 2004; Jeng et al., 2004), similar to ActA-motilty for *L. monocytogenes* (Welch et al., 1997, 1998; Loisel et al., 1999), IcsA-motility for *S. flexneri* (Egile et al., 1999; Loisel et al., 1999), and MirA-motility for *M. marinum* (Stamm et al., 2003, 2005; Hill and Welch, 2022). Thus, a consistent feature of Arp2/3-driven bacterial motility is that it continues following bacterial contact with the donor cell plasma membrane to generate long protrusions. In contrast, we observed bacteria at 1 dpi undergoing Sca2-motility ceased motility near the donor cell plasma membrane and spread into neighboring cells via short protrusions that exhibited reduced amplitudes of oscillation between elongation and retraction, and required less time to become engulfed by neighboring cells. Thus, the motility previously described for WT *R. parkeri* at 1 dpi (Lamason et al., 2016) appears to have been mediated by Sca2. These differences demonstrate that the two modes of *R. parkeri* motility result in two distinct modes of cell-to-cell spread during infection *in vitro*.

The observed differences in the mechanism and dynamics of RickA-spread versus Sca2-spread raises the question of whether there are distinctions with regards to the involvement of other bacterial and host factors. Actin-based motility and protrusion formation were shown to be sufficient for bacterial cell-to-cell spread by hijacking host processes of membrane exchange (Monack and Theriot, 2001). Studies of *L. monocytogenes* and *S. flexneri* have since revealed a variety of host proteins that are important for cell-to-cell spread. Protrusion formation during infection with *L. monocytogenes* and *S. flexneri* is mediated by host formins (Heindl et al., 2010; Fattouh et al., 2015) and the exocyst complex (Dowd et al., 2021; Herath et al., 2021). Stabilization of protrusions requires host plasma membrane proteins (Dhanda et al., 2019b) and proteins involved in linking the plasma membrane to the underlying actin cytoskeleton (Pust et al., 2005; Bishai et al., 2013; Dhanda et al., 2019a). Lastly, the engulfment of protrusions is mediated by host kinases (Chong et al., 2011; Dragoi and Agaisse, 2014, 2015), regulators of actin disassembly (Talman et al., 2014), and proteins involved in endocytosis (Fukumatsu et al., 2012; Sanderlin et al., 2019; Dhanda et al., 2020, 2021). Whether any of these host proteins are differentially involved in RickA-spread and/or Sca2-spread of *R. parkeri* remains to be elucidated. With regard to bacterial factors, *R. parkeri* utilizes Sca4 to interact with vinculin and disrupt its binding to α-catenin, manipulating intercellular tension and facilitating protrusion engulfment during Sca2-spread (Lamason et al., 2016). This is similar to manipulation of intercellular tension by *L. monocytogenes* using the effector internalin C (InlC) to disrupt Tuba/N-WASP complexes and dynamin 2 (Rajabian et al., 2009; Tijoriwalla et al., 2024), and by *S. flexneri* using IpaC to antagonize β-catenin (Shaikh et al., 2003; Duncan-Lowey et al., 2020), although these proteins are important for protrusion formation. Whether Sca4 or other bacterial factors are required for RickA-spread, perhaps by relieving intercellular tension for protrusion formation or resolution, remains unclear.

The differences between RickA-spread and Sca2-spread *in vitro* suggest that RickA and Sca2 may also play different roles *in vivo*. We previously showed that *sca2*::Tn *R. parkeri* strains are attenuated in lethality and dissemination from the skin to the liver and spleen following i.d. infection of *Ifnar1*^-/-^; *Ifngr1*^-/-^ DKO mice (Burke et al., 2021), and our current results extend these findings to the lung and brain. Attenuated dissemination to the spleen and liver was also reported following foodborne (Louie et al., 2019; Tucker et al., 2023) infection of animals with *L. monocytogenes* Δ*actA* mutants. How actin-based motility and cell-to-cell spread contribute to dissemination from the site of infection to other organs remains unclear. However, within the skin, we found that *sca2*::Tn bacteria formed smaller foci of infection than WT. Similarly, in the placenta of *L. monocytogenes*-infected pregnant guinea pigs (Bakardjiev et al., 2005) and mice (Le Monnier et al., 2007), and in the brain of neonatal mice (Pägelow et al., 2018), Δ*actA* bacteria form smaller foci of infection than WT. Furthermore, in colonic tissue of infant rabbits following rectal or oral infection with *S. flexneri*, Δ*icsA* bacteria form smaller foci of infection than WT (Yum et al., 2019; Kuehl et al., 2020). It is possible that actin-based motility and cell-to-cell spread within an organ facilitates spread to motile immune cells and the vasculature, facilitating further dissemination.

In contrast with the *sca2*::Tn mutant, the *rickA*::Tn mutant is not distinguishable from WT with regard to lethality or the extent of organ colonization. However, infection with *rickA*::Tn *R. parkeri* produces eschars that are significantly reduced in size and severity. *R. parkeri* mutants in genes encoding outer membrane protein B (*ompB*::Tn) (Burke et al., 2021) and protein lysine methyltransferases (PKMTs, *pkmt1*::Tn and *pkmt2*::Tn) (Engström et al., 2021) also cause smaller and less severe eschars following i.d. infection of *Ifnar1^-/-^*; *Ifngr1^-/-^* DKO mice. Nonetheless, in contrast to the *rickA*::Tn mutant, these other mutants are attenuated in lethality. Future studies comparing inflammation and infection in the skin between *ompB*::Tn, *pkmt1*::Tn, *pkmt2*::Tn, and *rickA*::Tn bacteria may provide insights into the specific contributions of bacterial growth, cell-to-cell spread, and inflammation to eschar formation.

Our results set the stage for future investigation into the function of RickA-driven and Sca2-driven actin-based motility and cell-to-cell spread at the cellular, tissue, and organismal scales. Our observations that RickA and Sca2 play differing roles *in vivo* raise the question of whether they may have distinct functions in spread between specialized cell-types or within specific tissues. Moreover, RickA-motility and Sca2-motility occur in tick cells, but neither factor is implicated in dissemination within the tick host (Harris et al., 2018). Whether they have roles in transmission to the mammalian host remains unclear. Further study of RickA and Sca2 will enhance our understanding of the evolution of different actin-based motility mechanisms and the roles of actin-based motility during infection.

## Materials and methods

### Mammalian cell culture

Frozen vials of human lung epithelial cells (A549), African green monkey kidney epithelial cells (Vero), and human embryonic kidney cells (HEK293T) were obtained from the UC Berkeley Cell Culture facility. All cells were grown at 37ºC in 5% CO_2_. Vials were thawed, centrifuged at 350 x g for 5 min, and passaged 3 times prior to use for experimentation under the conditions mentioned below. A549 and HEK293T cells were maintained in DMEM containing D-glucose (4.5 g/l) and L-glutamine (Gibco), supplemented with 10% heat-inactivated and filter-sterilized fetal bovine serum (ATLAS). Vero cells were maintained in DMEM containing D-glucose (4.5 g/l) and L-glutamine (Gibco), supplemented with 2% heat-inactivated and filter-sterilized fetal bovine serum (Gemcell).

### Generation of cell lines

To generate A549 cells expressing TagRFP-t-farnesyl, retroviral transduction was performed as previously outlined (Kostkow, N., Welch, MD., 2022) except 400 ng pCL-SIN-Ampho, 400 ng pCMV-VSVg, and 1200 ng pCLIP2B+TagRFPT-F (Naviaux et al., 1996; Lamason et al., 2016) were instead used to package viral particles. Additionally, polybrene was added directly into the filtered viral supernatant prior to transduction. Lastly, cells were selected using 1 μg/mL puromycin (Calbiochem, 540411) and cells were sorted using the Aria Fusion Cell Sorter (UC Berkeley Flow Cytometry Facility) for the top 50% brightest cells expressing TagRFP.

To generate A549 cells expressing TagRFP-t-farnesyl and Lifeact-3xTagBFP, lentiviruses were packaged in HEK293T cells plated 24 h prior as previously described (Lamason et al., 2016). Briefly, HEK293T cells were transfected via calcium phosphate transfection with 400 ng pMDLg-RRE, 400 ng pRSV-Rev, 400 ng pCMV-VSVg, and 800 ng pFCW2IB-Lifeact-3xTagBFP (Lamason et al., 2016). Similarly, cells were transduced as previously described (Kostow and Welch, 2022), except transduced cells were selected using 12 μg/mL blasticidin (Gibco, A1113903) and 1μg/mL puromycin (Lamason et al., 2016). Cells were sorted as described above for the top 50% brightest cells expressing TagRFP and top 0.1% brightest cells expressing TagBFP.

### Bacterial preparation

*R. parkeri* strain Portsmouth was obtained from the Centers for Disease Control and Prevention. The generation of the *rickA*::Tn and *sca2*::Tn mutants was described previously (Lamason et al., 2018). Bacteria were propagated by infecting multiple T175 flasks containing confluent monolayers of Vero cells with ∼1-5×10^6^ pfu *R. parkeri* per flask. For wild-type and *rickA*::Tn bacteria, infection progressed for ∼5 d, when cells of the monolayer would begin to round up and lift off the plate. For *sca2*::Tn bacteria, infection progressed for ∼6 d, when cells began rounding up and were beginning to lift off the plate. At each endpoint, infected cells were scraped and collected into 50 mL conical tubes. Tubes were then centrifuged at 4000 rpm for 15 min at 4ºC to pellet cells. The resulting cell pellets were resuspended in K-36 buffer (0.05 M KH_2_PO_4_, 0.05 M K_2_HPO_4_, 0.1 M KCl, 0.015 M NaCl, pH 7) and were Dounce homogenized for ∼40 strokes on ice. The suspension was then transferred to a 50 mL conical tube and was centrifuged at 200 x g for 5 min at 4ºC. The supernatant from each tube, which contained bacteria, was then overlayed on 30% RenoCal-76 (diluted in K-36 buffer; Bracco Diagnostics, 04H208) in ultracentrifuge tubes (Beckman/Coulter, 344058). Tubes were then placed in a SW-32TI rotor (Beckman/Coulter) and were centrifuged at 18,500 rpm for 20 min at 4ºC. The resulting bacterial pellet was resuspended in brain heart infusion (BHI) media (BD, 237500) to yield a “30% prep” which was stored at -80ºC. Titers for each “30% prep” bacterial stock were determined via plaque assay.

### Plaque assays

Monolayers of Vero cells plated on either 6- or 12-well plates were infected with bacteria (either from a 30% prep stock, from a homogenized organ, or from lysed cells) that were serially diluted in Vero cell media. Plates were then centrifuged at 300 x g for 5 min at room temperature, and infection progressed for at least 3 h at 33ºC. From 3-24 h post infection, media was aspirated and replaced with 4 or 2 mL agarose overlay (0.7% agarose in DMEM with 5% FBS) for 6- or 12-well plates, respectively. Infection progressed at 33ºC until plaques formed (∼5-7 d for wild-type and *rickA*::Tn bacteria, 7-14 d for *sca2*::Tn bacteria). When plaques were first visible, neutral red overlay (0.7% agarose in DMEM with 5% FBS and 2.5% neutral red, Sigma-Aldrich, N6264-50 mL) was applied to wells containing plaques. Plaques were then counted the following day.

To measure plaque size, 6-well plates containing plaques were scanned 24 h following addition of neutral red overlay. The scanned image was then analyzed using the ImageJ/FIJI (ver 1.0) (Schindelin et al., 2012) “Analyze Particles” function with the following settings: size = 0.2 – infinity; circularity = 0.3 – 1; and shows = outlines.

### Infectious focus size measurements

For infectious focus size assays, 2.5×10^5^ A549 cells were plated onto 12 mm^2^ glass coverslips (Fisherbrand, 12-545-81P) and were grown at 37ºC overnight. The following day, A549 cells were infected with wild-type or *rickA*::Tn bacteria at an MOI of 0.005 – 0.02. Bacteria were subsequently centrifuged at 300 x g for 5 min at room temperature and infection was allowed to progress at 33ºC. After 3 h, gentamycin was added to each well (final concentration was 10 μg/mL). Infection progressed at 33ºC for 2 d. Coverslips were then fixed with warm 4% paraformaldehyde (Ted Pella Inc., 18505) for 10 min, washed with PBS, and stained for *Rickettsia* using mouse anti-*Rickettsia* 14-13 (1:1000; gift from T. Hackstadt, Anacker, R.L., *et al*. 1987) and DAPI (1:300; Invitrogen, D1306). Coverslips were imaged on an Olympus IX71 inverted microscope equipped with an Olympus LUCPlanFL N 20x/0.45 objective and optiMOS camera (QImaging). Images were acquired using µManager software (ver 1.4.20) (Edelstein et al., 2010, 2014). Images were taken of 10 infectious foci per strain per experiment. Focus sizes were quantified using a customized CellProfiler pipeline (ver 4.0.4) (Stirling et al., 2021) that identified nuclei based on DAPI intensity and size. Cell boundaries were approximated using a fixed distance of 50 pixels from each nuclei, and bacteria were identified based on anti-*Rickettsia* 14-13 staining intensity and size. These data were then used to relate bacteria to cell boundaries to identify infected and uninfected cells. The number of infected cells per focus was then recorded in Prism 10 version 10.2.3 for data visualization and statistics (Sanderlin et al., 2019).

### Live cell imaging of cell-cell spread

To visualize RickA-mediated cell-to-cell spread using *sca2*::Tn mutant bacteria, A549 cells were plated onto 35 mm MatTek dishes (MatTek Corporation, P35G-1.5.20-C), and cells were grown overnight at 37ºC. For 3 of 10 videos, 12×10^5^ A549 cells expressing only TagRFP-t-farnesyl were plated onto MatTek dishes and were transiently transfected the following day via Lipofectamine 2000 (ThermoFisher Scientific, 11668027) and 1 μg pFCW2IB-Lifeact-3xTagBFP per chamber. The following day, transfected A549 cells were infected at an MOI of 10 with *sca2*::Tn bacteria diluted in Ringer’s buffer (155 mM NaCl, 5 mM KCl, 2 mM CaCl_2_, 1 mM MgCl_2_, 2 mM NaH_2_PO_4_, 50 mM HEPES, 10 mM glucose) with 10% FBS, 1:100 oxyrase (VWR, 101975-866), and 10 mM succinate. For the remaining 7 of 10 videos, 8×10^5^ A549 cells stably expressing TagRFP-t-farnesyl and Lifeact-3xTagBFP (described above) were plated onto 33 mm MatTek dishes grown for 2 d at 37ºC. Cells were then infected with *sca2*::Tn bacteria diluted in Ringer’s buffer with 10% FBS, 1:100 oxyrase, and 10mM succinate by centrifuging bacteria onto cells at 300 x g for 5 min at room temperature. Immediately afterwards, infected cells were imaged using a Nikon Ti Eclipse microscope with a 60X (1.4 NA) Plan Apo objective, a Clara Interline CCD Camera, a Yokogawa CSU-XI spinning disc confocal, in an environmental chamber set at 33ºC (Lamason et al., 2016). Images were taken at 20 s intervals for times ranging from 45 min to 2 h.

To visualize Sca2-mediated cell-to-cell spread using *rickA*::Tn bacteria, 8×10^5^ A549 cells expressing TagRFP-t-farnesyl and Lifeact-3xTagBFP (described above) were plated onto 33 mm MatTek dishes, and cells were grown overnight at 37ºC. The following day, *rickA*::Tn bacteria were diluted in A549 cell media and were used to infect A549 cells at an MOI of 3. Cells were infected by centrifuging bacteria onto cells at 300 x g for 5 min at room temperature. Infection was allowed to progress at 33ºC for 27 h, at which point A549 cell media was replaced with Ringer’s buffer with 10% FBS, 1:100 oxyrase, and 10 mM succinate and cells were imaged as described above.

To quantify the kinetics of RickA- and Sca2-mediated cell-to-cell spread, videos and stills were analyzed using FIJI/ImageJ version 1.0. To measure maximum protrusion lengths, still images were collected from videos of successful cell-cell spread events. Protrusion length was measured by tracing the protrusion from its base (at the junction with the rest of the host cell plasma membrane) to its tip (at the distal tip of the bacterium) with three technical replicates, and then averaging the three measurements to obtain the reported “maximum length” of the protrusion. To measure changes to protrusion length over time, single particle tracking of the protrusion distal tip was performed. This analysis resulted in a dataset of X- and Y-coordinates from the protrusion base (which served as the “origin”, (0, 0)) at each 20 s interval. These coordinates were then used to calculate the direct distance from the protrusion tip to the protrusion base over time. To quantify the distance traveled by a spreading bacterium between each 20 s interval and its directionality over time, we subtracted the distance between the protrusion tip and the protrusion base (previously determined when calculating changes in protrusion length) at T_n_ and T_n-1_. The value of the absolute length of fluctuations by a bacterium spreading was calculated by measuring the direct distance between the protrusion tips at T_n_ and T_n-1_, which does not include information on directionality, but includes the absolute value of the distance traveled between each consecutive frame. These distances were added to yield the total distance traveled by a bacterium during a cell-to-cell spread event. Lastly, the time from initiating cell-to-cell spread to its resolution was determined by measuring the time in between the formation of a protrusion (when a bacterium makes an indent on the plasma membrane) to its resolution in a recipient cell (when a bacterium is clearly in a secondary vacuole).

### Mouse studies

Animal research was conducted under a protocol approved by the University of California, Berkeley, Institutional Animal Care and Use Committee (IACUC) in compliance with the Animal Welfare Act and other federal statutes relating to animals and experiments using animals (Welch lab animal use protocol AUP-2016-09-8426-2). All mice were of the C57BL/6J background and the *Ifnar1^-/-^; Ifngr1^-/-^* DKO genotype carrying mutations in the genes encoding the receptors for IFN-I (*Ifnar1*) and IFN-γ (*Ifngr1*) (Jackson Labs stock # 029098). Mice were between 8-20 weeks old and were healthy at the time of initial infection. Mice were selected for experiments based on their availability, regardless of sex, and both sexes were used for each experimental group. For infections, *R. parkeri* was prepared by diluting 30% prep stocks in cold, sterile PBS on ice, to the desired concentration (2×10^4^ pfu/ml for intradermal infection) and were kept on ice during injections. Mice were anesthetized with 3-4% isoflurane via inhalation. The right flank of each mouse was shaved with a hair trimmer (BrainTree Scientific Inc, CLP-41590P) and the skin was wiped with 70% ethanol. 50 µl of bacterial suspension in PBS was injected intradermally using a 30-gauge needle. Mice were monitored until they were fully awake. No adverse effects were recorded from the anesthesia. A small aliquot of diluted bacteria was also set aside for a plaque assay to confirm the number of viable bacteria in the suspension.

All mice in this study were monitored every other day for clinical signs of disease throughout the course of infection. Body temperature was monitored using a rodent rectal thermometer (BrainTree Scientific Inc, RET-3). Mice were euthanized using CO_2_ followed by cervical dislocation upon presentation of core body temperatures less than 90ºF, and/or severe lethargy, paralysis, facial edema, or necrotic tails. Mice presenting with scruffed fur and mild-moderate lethargy were monitored daily for recovery or were euthanized if symptoms did not improve after 14 d. Mice presenting with eschars at the site of infection but lacking other signs of serious infection were kept alive and monitored every other day until a pre-determined endpoint.

To measure the bacterial burden in various organs, mice were euthanized at pre-determined endpoints and doused with ethanol. Mouse organs were extracted and divided in half. One half of each organ was deposited into a 50 mL conical tube containing 30-40 mL 10% neutral buffered formalin (Sigma, HT501128) for histology studies (explained in the following section). The other half was deposited into a pre-weighed 50 mL conical tube containing 2 mL (brain), 4 mL (for spleen, lung, skin), or 8 mL PBS (liver). These tubes were re-weighed after the organs were deposited to determine the weight of the organ itself. Organs were kept on ice and were homogenized for ∼10 s using an immersion homogenizer at ∼23,000 rpm. Organ homogenates were spun down at 260 x g for 5 min at 4ºC to pellet tissue debris, and freed bacteria in the supernatant were enumerated in plaque assays (described above) to quantify the bacterial burden in each organ. After about 1 h of infection, carbenicillin and amphotericin B were diluted into each well to a final concentration of 50 μg/mL and 1 μg/mL, respectively. The following day, cells were gently washed by replacing the existing media with 500 μL DMEM containing 2% FBS. The media was then aspirated and replaced with 2 mL agarose overlay (0.7% agarose, 5% FBS, and 1μg/mL amphotericin B). When plaques were visible (4-14 d post infection), 1 mL of agarose overlay containing neutral red (0.7% agarose, 5% FBS, 1 μg/mL amphotericin B, and 2.5% neutral red) was added to each well. Plaques were counted the following day. In some scenarios, plaque assays for a specific mouse’s tissue, such as the spleen, were contaminated and therefore were not included in the final dataset while results from other plaque assays involving other organs from the same mouse were included.

### Histology

For histology experiments, half-organs extracted from mice as described above were fixed in 10% buffered formalin, embedded in paraffin, sectioned, and stained with hematoxylin and eosin (H&E). Additional serial sections of all tissues were submitted to indirect immunohistochemistry (IHC) for the presence of *Rickettsia* using the rabbit anti-*Rickettsia* I7205 antibody (1:500 dilution; gift from Ted Hackstadt). Histology was performed by HistoWiz Inc. (histowiz.com) using a standard operating procedure and fully automated workflow. Samples were processed, embedded in paraffin, and sectioned at 4 μm. Immunohistochemistry was performed on a Bond Rx autostainer (Leica Biosystems) using standard protocols. Bond Polymer Refine Detection (Leica Biosystems) was used according to the manufacturer’s protocol. After staining, sections were dehydrated and film coverslipped using a TissueTek-Prisma and Coverslipper (Sakura). Whole slide scanning (40x) was performed on an Aperio AT2 (Leica Biosystems). Blinded histologic evaluation and semiquantitative scoring were blindly performed by a board-certified pathologist. Tissue inflammation was scored as follows: 0, absent (tissue is unremarkable); 1, minimal (rare finding); 2, mild (finding is more noticeable, affecting <25% of the tissue); 3, moderate (finding is prominent, affecting >25% and <50% of the tissue); 4, marked (finding is striking, affecting >50% of the tissue). And immunohistochemical labeling was scored as follows: 0, none; 1, sparse foci with few bacteria; 2, multiple foci with several bacteria; 3, coalescing foci with numerous bacteria; 4, abundant bacteria throughout the entire tissue. Links to histological images and immunohistochemistry data are available upon requests to the authors.

### Statistical analyses

The statistical analyses and significance are reported in the figure legends. Data were considered to be statistically significant when p-values were less than 0.05, as determined by mixed-effects analysis, two-tailed Mann-Whitney tests, Dunn’s multiple comparison test, or by log-rank (Mantel-Cox) test. Error bars indicated standard deviation for all experiments. Statistical analyses were performed using GraphPad PRISM version 10.2.3.

## Acknowledgements

We thank Ted Hackstadt and Christopher Paddock for kindly providing strains and reagents. We thank Rebecca Lamason, Shawna Reed, and Thomas Burke, whose work set the stage for this project; as well as other past and current members of the Welch Lab for their suggestions throughout the project. We also thank Thomas Burke, Victoria Chevée, Rafael Rivera-Lugo, Jesse Garcia Castillo, and the UC Berkeley animal facility staff for training Cuong Joseph Tran and Zahra Zubair-Nizami to work with mice. We thank the following UC Berkeley core facilities and their staff for providing equipment, reagents, and technical support to complete this work: Kartoosh Heydari, Anita Lin, Melaine Delcroix, and Harman Dhaliwal (CRL Flow Cytometry Core); Holly Aaron and Feather Ives (CRL Molecular Imaging Center); and Alison Killilea (Cell Culture Facility). We thank Daniel Portnoy, Sarah Stanley, and Kathleen Ryan for their technical discussion and guidance during this work. Lastly, we thank Neil Fischer for proofreading the manuscript. This work was funded by NIH/NIAID grant R01 AI109044 to M.D.W. C.J.T. was funded by the Berkeley Fellowship for Graduate Study; the Schwartz-Efrusy-Levy Global Health Doctoral Fellowship; the Eki & Nobuta Akahoshi and Seiko Baba Brodbeck Endowed Fund; the Irving H. Wiesenfeld Fellowship; and the UC Dissertation-Year Fellowship.

## References

Alqassim, S. S., Lee, I.-G., and Dominguez, R. (2019). *Rickettsia* Sca2 Recruits Two Actin Subunits for Nucleation but Lacks WH2 Domains. Biophysical Journal 116, 540–550. doi: 10.1016/j.bpj.2018.12.009

Bakardjiev, A. I., Stacy, B. A., and Portnoy, D. A. (2005). Growth of *Listeria monocytogenes* in the guinea pig placenta and role of cell-to-cell spread in fetal infection. J Infect Dis 191, 1889–1897. doi: 10.1086/430090

Bernardini, M. L., Mounier, J., d’Hauteville, H., Coquis-Rondon, M., and Sansonetti, P. J. (1989). Identification of icsA, a plasmid locus of *Shigella flexneri* that governs bacterial intra- and intercellular spread through interaction with F-actin. Proceedings of the National Academy of Sciences 86, 3867–3871. doi: 10.1073/pnas.86.10.3867

Bishai, E. A., Sidhu, G. S., Li, W., Dhillon, J., Bohil, A. B., Cheney, R. E., et al. (2013). Myosin-X facilitates *Shigella*-induced membrane protrusions and cell-to-cell spread. Cellular Microbiology 15, 353–367. doi: 10.1111/cmi.12051

Burke, T. P., Engström, P., Tran, C. J., Langohr, I. M., Glasner, D. R., Espinosa, D. A., et al. (2021). Interferon receptor-deficient mice are susceptible to eschar-associated rickettsiosis. eLife 10, e67029. doi: 10.7554/eLife.67029

Campellone, K. G., and Welch, M. D. (2010). A nucleator arms race: cellular control of actin assembly. Nat Rev Mol Cell Biol 11, 237–251. doi: 10.1038/nrm2867

Chong, R., Squires, R., Swiss, R., and Agaisse, H. (2011). RNAi Screen Reveals Host Cell Kinases Specifically Involved in *Listeria monocytogenes* Spread from Cell to Cell. PLOS ONE 6, e23399. doi: 10.1371/journal.pone.0023399

Dantas-Torres, F. (2007). Rocky Mountain spotted fever. The Lancet Infectious Diseases 7, 724–732. doi: 10.1016/S1473-3099(07)70261-X

Dhanda, A. S., Lulic, K. T., Vogl, A. W., Mc Gee, M. M., Chiu, R. H., and Guttman, J. A. (2019a). *Listeria Membrane* Protrusion Collapse: Requirement of Cyclophilin A for Listeria Cell-to-Cell Spreading. J Infect Dis 219, 145–153. doi: 10.1093/infdis/jiy255

Dhanda, A. S., Lulic, K. T., Yu, C., Chiu, R. H., Bukrinsky, M., and Guttman, J. A. (2019b). *Listeria monocytogenes* hijacks CD147 to ensure proper membrane protrusion formation and efficient bacterial dissemination. Cell. Mol. Life Sci. 76, 4165–4178. doi: 10.1007/s00018-019-03130-4

Dhanda, A. S., Vogl, A. W., Ness, F., Innocenti, M., and Guttman, J. A. (2021). mDia1 Assembles a Linear F-Actin Coat at Membrane Invaginations To Drive *Listeria monocytogenes* Cell-to-Cell Spreading. mBio 12, e0293921. doi: 10.1128/mBio.02939-21

Dhanda, A. S., Yu, C., Lulic, K. T., Vogl, A. W., Rausch, V., Yang, D., et al. (2020). *Listeria monocytogenes* Exploits Host Caveolin for Cell-to-Cell Spreading. mBio 11, 10.1128/mbio.02857-19. doi: 10.1128/mbio.02857-19

Domann, E., Wehland, J., Rohde, M., Pistor, S., Hartl, M., Goebel, W., et al. (1992). A novel bacterial virulence gene in *Listeria monocytogenes* required for host cell microfilament interaction with homology to the proline-rich region of vinculin. EMBO J 11, 1981–1990. doi: 10.1002/j.1460-2075.1992.tb05252.x

Dowd, G. C., Mortuza, R., and Ireton, K. (2021). Molecular Mechanisms of Intercellular Dissemination of Bacterial Pathogens. Trends in Microbiology 29, 127–141. doi: 10.1016/j.tim.2020.06.008

Dragoi, A.-M., and Agaisse, H. (2014). The Serine/Threonine Kinase STK11 Promotes *Shigella flexneri* Dissemination through Establishment of Cell-Cell Contacts Competent for Tyrosine Kinase Signaling. Infect Immun 82, 4447–4457. doi: 10.1128/IAI.02078-14

Dragoi, A.-M., and Agaisse, H. (2015). The class II phosphatidylinositol 3-phosphate kinase PIK3C2A promotes *Shigella flexneri* dissemination through formation of vacuole-like protrusions. Infect Immun 83, 1695–1704. doi: 10.1128/IAI.03138-14

Duncan-Lowey, J. K., Wiscovitch, A. L., Wood, T. E., Goldberg, M. B., and Russo, B. C. (2020). *Shigella flexneri* Disruption of Cellular Tension Promotes Intercellular Spread. Cell Rep 33, 108409. doi: 10.1016/j.celrep.2020.108409

Edelstein, A., Amodaj, N., Hoover, K., Vale, R., and Stuurman, N. (2010). Computer control of microscopes using µManager. Curr Protoc Mol Biol Chapter 14, Unit14.20. doi: 10.1002/0471142727.mb1420s92

Edelstein, A. D., Tsuchida, M. A., Amodaj, N., Pinkard, H., Vale, R. D., and Stuurman, N. (2014). Advanced methods of microscope control using μManager software. J Biol Methods 1, e10. doi: 10.14440/jbm.2014.36

Egile, C., Loisel, T. P., Laurent, V., Li, R., Pantaloni, D., Sansonetti, P. J., et al. (1999). Activation of the CDC42 effector N-WASP by the *Shigella flexneri* IcsA protein promotes actin nucleation by Arp2/3 complex and bacterial actin-based motility. J Cell Biol 146, 1319–1332. doi: 10.1083/jcb.146.6.1319

El Karkouri, K., Ghigo, E., Raoult, D., and Fournier, P.-E. (2022). Genomic evolution and adaptation of arthropod-associated *Rickettsia*. Sci Rep 12, 3807. doi: 10.1038/s41598-022-07725-z

Engström, P., Burke, T. P., Tran, C. J., Iavarone, A. T., and Welch, M. D. (2021). Lysine methylation shields an intracellular pathogen from ubiquitylation and autophagy. Science Advances 7, eabg2517. doi: 10.1126/sciadv.abg2517

Fattouh, R., Kwon, H., Czuczman, M. A., Copeland, J. W., Pelletier, L., Quinlan, M. E., et al. (2015). The diaphanous-related formins promote protrusion formation and cell-to-cell spread of *Listeria monocytogenes*. J Infect Dis 211, 1185–1195. doi: 10.1093/infdis/jiu546

Fukumatsu, M., Ogawa, M., Arakawa, S., Suzuki, M., Nakayama, K., Shimizu, S., et al. (2012). Shigella Targets Epithelial Tricellular Junctions and Uses a Noncanonical Clathrin-Dependent Endocytic Pathway to Spread Between Cells. Cell Host & Microbe 11, 325–336. doi: 10.1016/j.chom.2012.03.001

Gillespie, J. J., Beier, M. S., Rahman, M. S., Ammerman, N. C., Shallom, J. M., Purkayastha, A., et al. (2007). Plasmids and Rickettsial Evolution: Insight from *Rickettsia felis*. PLOS ONE 2, e266. doi: 10.1371/journal.pone.0000266

Gouin, E., Egile, C., Dehoux, P., Villiers, V., Adams, J., Gertler, F., et al. (2004). The RickA protein of *Rickettsia conorii* activates the Arp2/3 complex. Nature 427, 457–461. doi: 10.1038/nature02318

Gouin, E., Gantelet, H., Egile, C., Lasa, I., Ohayon, H., Villiers, V., et al. (1999). A comparative study of the actin-based motilities of the pathogenic bacteria *Listeria monocytogenes*, *Shigella flexneri* and *Rickettsia conorii*. J Cell Sci 112 (Pt 11), 1697–1708. doi: 10.1242/jcs.112.11.1697

Haglund, C. M., Choe, J. E., Skau, C. T., Kovar, D. R., and Welch, M. D. (2010). *Rickettsia* Sca2 is a bacterial formin-like mediator of actin-based motility. Nat Cell Biol 12, 1057– 1063. doi: 10.1038/ncb2109

Harris, E. K., Jirakanwisal, K., Verhoeve, V. I., Fongsaran, C., Suwanbongkot, C., Welch, M. D., et al. (2018). Role of Sca2 and RickA in the Dissemination of *Rickettsia parkeri* in *Amblyomma maculatum*. Infection and Immunity 86, 10.1128/iai.00123-18. doi: 10.1128/iai.00123-18

Heindl, J. E., Saran, I., Yi, C., Lesser, C. F., and Goldberg, M. B. (2010). Requirement for Formin-Induced Actin Polymerization during Spread of *Shigella flexneri*. Infection and Immunity 78, 193–203. doi: 10.1128/iai.00252-09

Herath, T. U. B., Roy, A., Gianfelice, A., and Ireton, K. (2021). *Shigella flexneri* subverts host polarized exocytosis to enhance cell-to-cell spread. Molecular Microbiology 116, 1328– 1346. doi: 10.1111/mmi.14827

Hill, N. S., and Welch, M. D. (2022). A glycine-rich PE_PGRS protein governs mycobacterial actin-based motility. Nat Commun 13, 3608. doi: 10.1038/s41467-022-31333-0

Jeng, R. L., Goley, E. D., D’Alessio, J. A., Chaga, O. Y., Svitkina, T. M., Borisy, G. G., et al. (2004). A *Rickettsia* WASP-like protein activates the Arp2/3 complex and mediates actin-based motility. Cellular Microbiology 6, 761–769. doi: 10.1111/j.1462-5822.2004.00402.x

Kadurugamuwa, J. L., Rohde, M., Wehland, J., and Timmis, K. N. (1991). Intercellular spread of *Shigella flexneri* through a monolayer mediated by membranous protrusions and associated with reorganization of the cytoskeletal protein vinculin. Infection and Immunity 59, 3463–3471. doi: 10.1128/iai.59.10.3463-3471.1991

Kleba, B., Clark, T. R., Lutter, E. I., Ellison, D. W., and Hackstadt, T. (2010). Disruption of the *Rickettsia rickettsii Sca2* Autotransporter Inhibits Actin-Based Motility. Infection and Immunity 78, 2240–2247. doi: 10.1128/iai.00100-10

Kocks, C., Gouin, E., Tabouret, M., Berche, P., Ohayon, H., and Cossart, P. (1992). *L. monocytogenes*-induced actin assembly requires the actA gene product, a surface protein. Cell 68, 521–531. doi: 10.1016/0092-8674(92)90188-i

Kostow, N., and Welch, M. D. (2022). Plasma membrane protrusions mediate host cell–cell fusion induced by *Burkholderia thailandensis*. MBoC 33, ar70. doi: 10.1091/mbc.E22-02-0056

Kuehl, C. J., D’Gama, J. D., Warr, A. R., and Waldor, M. K. (2020). An Oral Inoculation Infant Rabbit Model for *Shigella* Infection. mBio 11, 10.1128/mbio.03105-19. doi: 10.1128/mbio.03105-19

Lamason, R. L., Bastounis, E., Kafai, N. M., Serrano, R., del Álamo, J. C., Theriot, J. A., et al. (2016). *Rickettsia* Sca4 Reduces Vinculin-Mediated Intercellular Tension to Promote Spread. Cell 167, 670–683.e10. doi: 10.1016/j.cell.2016.09.023

Lamason, R. L., Kafai, N. M., and Welch, M. D. (2018). A streamlined method for transposon mutagenesis of *Rickettsia parkeri* yields numerous mutations that impact infection. PLOS ONE 13, e0197012. doi: 10.1371/journal.pone.0197012

Lamason, R. L., and Welch, M. D. (2017). Actin-based motility and cell-to-cell spread of bacterial pathogens. Current Opinion in Microbiology 35, 48–57. doi: 10.1016/j.mib.2016.11.007

Le Monnier, A., Autret, N., Join-Lambert, O. F., Jaubert, F., Charbit, A., Berche, P., et al. (2007). ActA Is Required for Crossing of the Fetoplacental Barrier by *Listeria monocytogenes*. Infection and Immunity 75, 950–957. doi: 10.1128/iai.01570-06

Loisel, T. P., Boujemaa, R., Pantaloni, D., and Carlier, M. F. (1999). Reconstitution of actin-based motility of *Listeria* and *Shigella* using pure proteins. Nature 401, 613–616. doi: 10.1038/44183

Louie, A., Zhang, T., Becattini, S., Waldor, M. K., and Portnoy, D. A. (2019). A Multiorgan Trafficking Circuit Provides Purifying Selection of *Listeria monocytogenes* Virulence Genes. mBio 10, 10.1128/mbio.02948-19. doi: 10.1128/mbio.02948-19

Madasu, Y., Suarez, C., Kast, D. J., Kovar, D. R., and Dominguez, R. (2013). *Rickettsia* Sca2 has evolved formin-like activity through a different molecular mechanism. Proceedings of the National Academy of Sciences 110, E2677–E2686. doi: 10.1073/pnas.1307235110

Monack, D. M., and Theriot, J. A. (2001). Actin-based motility is sufficient for bacterial membrane protrusion formation and host cell uptake. Cellular Microbiology 3, 633–647. doi: 10.1046/j.1462-5822.2001.00143.x

Naviaux, R. K., Costanzi, E., Haas, M., and Verma, I. M. (1996). The pCL vector system: rapid production of helper-free, high-titer, recombinant retroviruses. J Virol 70, 5701–5705. doi: 10.1128/JVI.70.8.5701-5705.1996

Paddock, C. D., Sumner, J. W., Comer, J. A., Zaki, S. R., Goldsmith, C. S., Goddard, J., et al. (2004). *Rickettsia parkeri*: A Newly Recognized Cause of Spotted Fever Rickettsiosis in the United States. Clinical Infectious Diseases 38, 805–811. doi: 10.1086/381894

Pägelow, D., Chhatbar, C., Beineke, A., Liu, X., Nerlich, A., van Vorst, K., et al. (2018). The olfactory epithelium as a port of entry in neonatal neurolisteriosis. Nat Commun 9, 4269. doi: 10.1038/s41467-018-06668-2

Pust, S., Morrison, H., Wehland, J., Sechi, A. S., and Herrlich, P. (2005). *Listeria monocytogenes* exploits ERM protein functions to efficiently spread from cell to cell. The EMBO Journal 24, 1287–1300. doi: 10.1038/sj.emboj.7600595

Rajabian, T., Gavicherla, B., Heisig, M., Müller-Altrock, S., Goebel, W., Gray-Owen, S. D., et al. (2009). The bacterial virulence factor InlC perturbs apical cell junctions and promotes cell-to-cell spread of *Listeria*. Nat Cell Biol 11, 1212–1218. doi: 10.1038/ncb1964

Reed, S. C. O., Lamason, R. L., Risca, V. I., Abernathy, E., and Welch, M. D. (2014). *Rickettsia* Actin-Based Motility Occurs in Distinct Phases Mediated by Different Actin Nucleators. Current Biology 24, 98–103. doi: 10.1016/j.cub.2013.11.025

Robbins, J. R., Barth, A. I., Marquis, H., de Hostos, E. L., Nelson, W. J., and Theriot, J. A. (1999). *Listeria monocytogenes* Exploits Normal Host Cell Processes to Spread from Cell to Cell. Journal of Cell Biology 146, 1333–1350. doi: 10.1083/jcb.146.6.1333

Sanderlin, A. G., Vondrak, C., Scricco, A. J., Fedrigo, I., Ahyong, V., and Lamason, R. L. (2019). RNAi screen reveals a role for PACSIN2 and caveolins during bacterial cell-to-cell spread. MBoC 30, 2124–2133. doi: 10.1091/mbc.E19-04-0197

Schindelin, J., Arganda-Carreras, I., Frise, E., Kaynig, V., Longair, M., Pietzsch, T., et al. (2012). Fiji: an open-source platform for biological-image analysis. Nat Methods 9, 676–682. doi: 10.1038/nmeth.2019

Scott, A. T., Vondrak, C. J., Sanderlin, A. G., and Lamason, R. L. (2022). Rickettsia parkeri. Trends in Microbiology 30, 511–512. doi: 10.1016/j.tim.2022.01.001

Sechi, A. S., Wehland, J., and Small, J. V. (1997). The Isolated Comet Tail Pseudopodium of *Listeria monocytogenes*: A Tail of Two Actin Filament Populations, Long and Axial and Short and Random. Journal of Cell Biology 137, 155–167. doi: 10.1083/jcb.137.1.155

Shaikh, N., Terajima, J., and Watanabe, H. (2003). IpaC of *Shigella* binds to the C-terminal domain of β-catenin. Microbial Pathogenesis 35, 107–117. doi: 10.1016/S0882-4010(03)00093-7

Stamm, L. M., Morisaki, J. H., Gao, L.-Y., Jeng, R. L., McDonald, K. L., Roth, R., et al. (2003). *Mycobacterium marinum* Escapes from Phagosomes and Is Propelled by Actin-based Motility. J Exp Med 198, 1361–1368. doi: 10.1084/jem.20031072

Stamm, L. M., Pak, M. A., Morisaki, J. H., Snapper, S. B., Rottner, K., Lommel, S., et al. (2005). Role of the WASP family proteins for *Mycobacterium marinum* actin tail formation. Proc Natl Acad Sci U S A 102, 14837–14842. doi: 10.1073/pnas.0504663102

Stirling, D. R., Swain-Bowden, M. J., Lucas, A. M., Carpenter, A. E., Cimini, B. A., and Goodman, A. (2021). CellProfiler 4: improvements in speed, utility and usability. BMC Bioinformatics 22, 433. doi: 10.1186/s12859-021-04344-9

Talman, A. M., Chong, R., Chia, J., Svitkina, T., and Agaisse, H. (2014). Actin network disassembly powers dissemination of *Listeria monocytogenes*. Journal of Cell Science 127, 240–249. doi: 10.1242/jcs.140038

Tijoriwalla, S., Liyanage, T., Herath, T. U. B., Lee, N., Rehman, A., Gianfelice, A., et al. (2024). The host GTPase Dynamin 2 modulates apical junction structure to control cell-to-cell spread of *Listeria monocytogenes*. Infect Immun, e0013624. doi: 10.1128/iai.00136-24

Tucker, J. S., Cho, J., Albrecht, T. M., Ferrell, J. L., and D’Orazio, S. E. F. (2023). Egress of *Listeria monocytogenes* from Mesenteric Lymph Nodes Depends on Intracellular Replication and Cell-to-Cell Spread. Infection and Immunity 91, e00064–23. doi: 10.1128/iai.00064-23

Welch, M. D., Iwamatsu, A., and Mitchison, T. J. (1997). Actin polymerization is induced by Arp2/3 protein complex at the surface of *Listeria monocytogenes*. Nature 385, 265–269. doi: 10.1038/385265a0

Welch, M. D., Rosenblatt, J., Skoble, J., Portnoy, D. A., and Mitchison, T. J. (1998). Interaction of Human Arp2/3 Complex and the *Listeria monocytogenes* ActA Protein in Actin Filament Nucleation. Science 281, 105–108. doi: 10.1126/science.281.5373.105

Yum, L. K., Byndloss, M. X., Feldman, S. H., and Agaisse, H. (2019). Critical role of bacterial dissemination in an infant rabbit model of bacillary dysentery. Nat Commun 10, 1826. doi: 10.1038/s41467-019-09808-4

