## Supplementary Figures S1-S3 for "The *Rickettsia* actin-based motility effectors RickA and Sca2 contribute differently to cell-to-cell spread and pathogenicity"

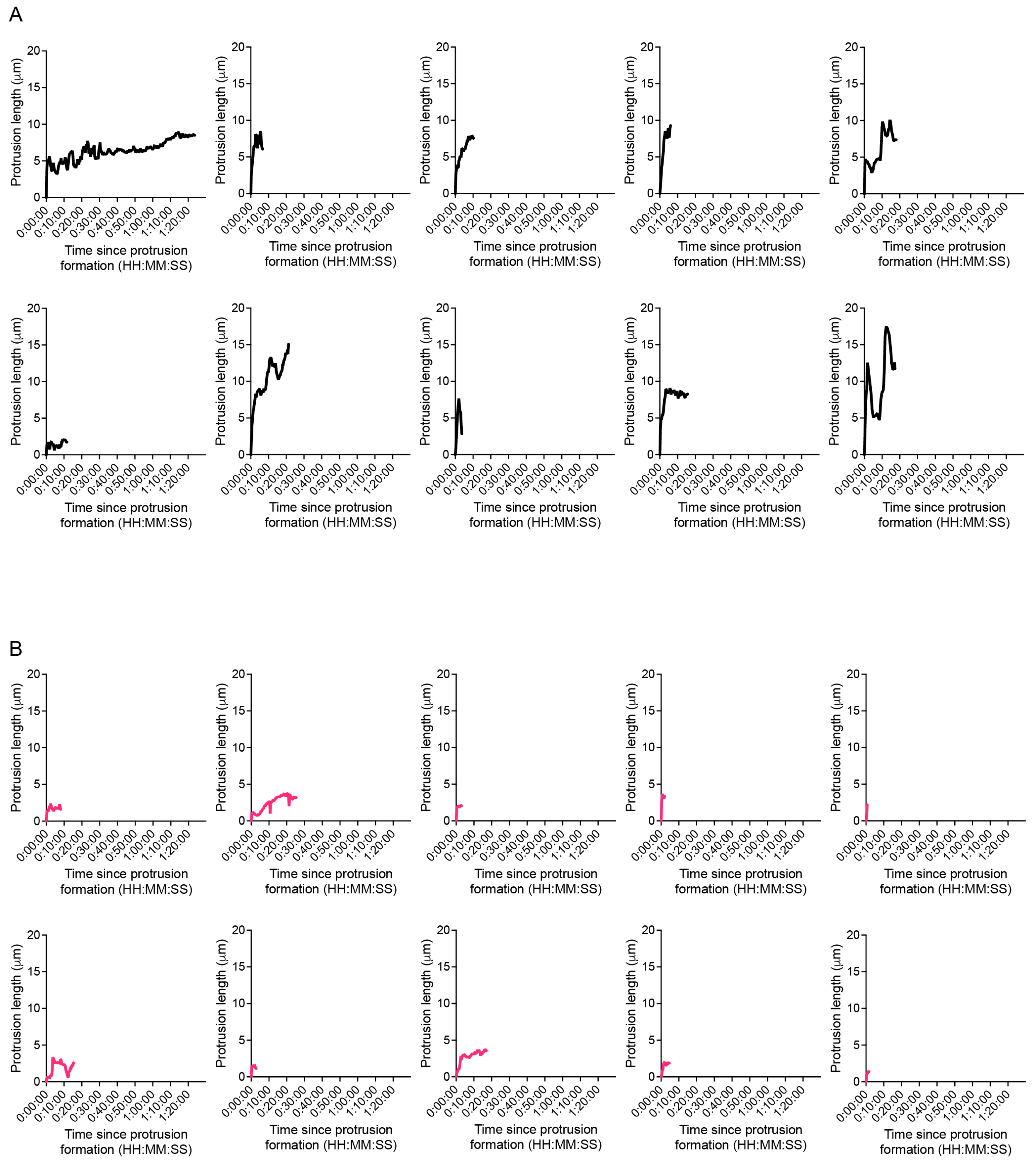


**Figure S1, related to Figure 4 – Protrusion lengths during individual spread events.**

Each graph shows the length of an individual protrusion over time during **(A)** RickA-spread (*sca2*::Tn mutant) or **(B)** Sca2-spread (*rickA*::Tn mutant). These data are included in Figure 4b.

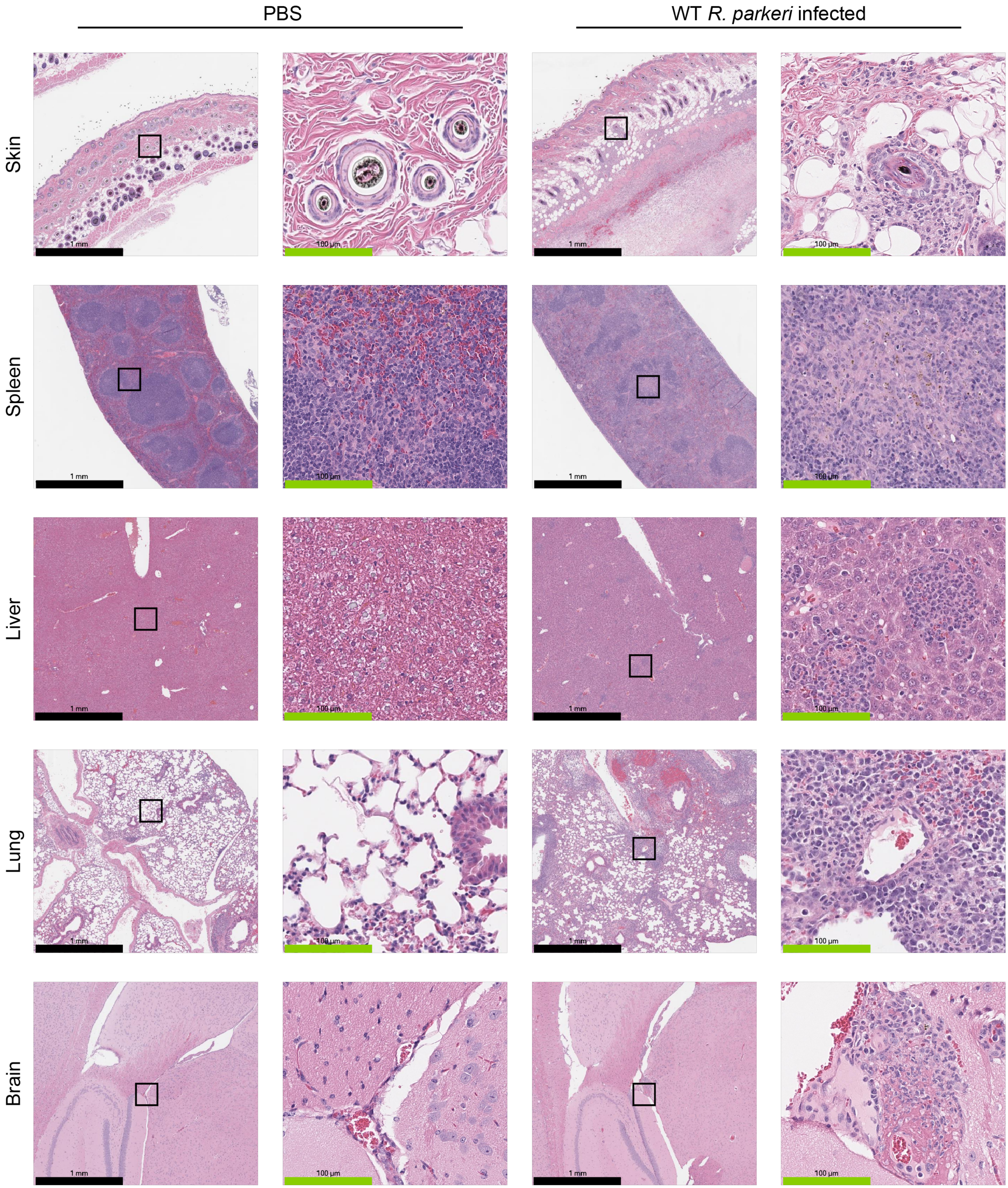


**Figure S2, related to Figure 7 – H&E staining of various organs.**

Representative images of H&E staining of the indicated organs in mice either injected i.d. with PBS (control) or infected with $10^3$ pfu WT *R. parkeri*. Results were qualitatively similar for *rickA*::Tn-infected and *sca2*::Tn-infected mice. Black squares on the left were magnified on the right. Black scale bars represent 1 mm, green scale bars represent 100 μm.

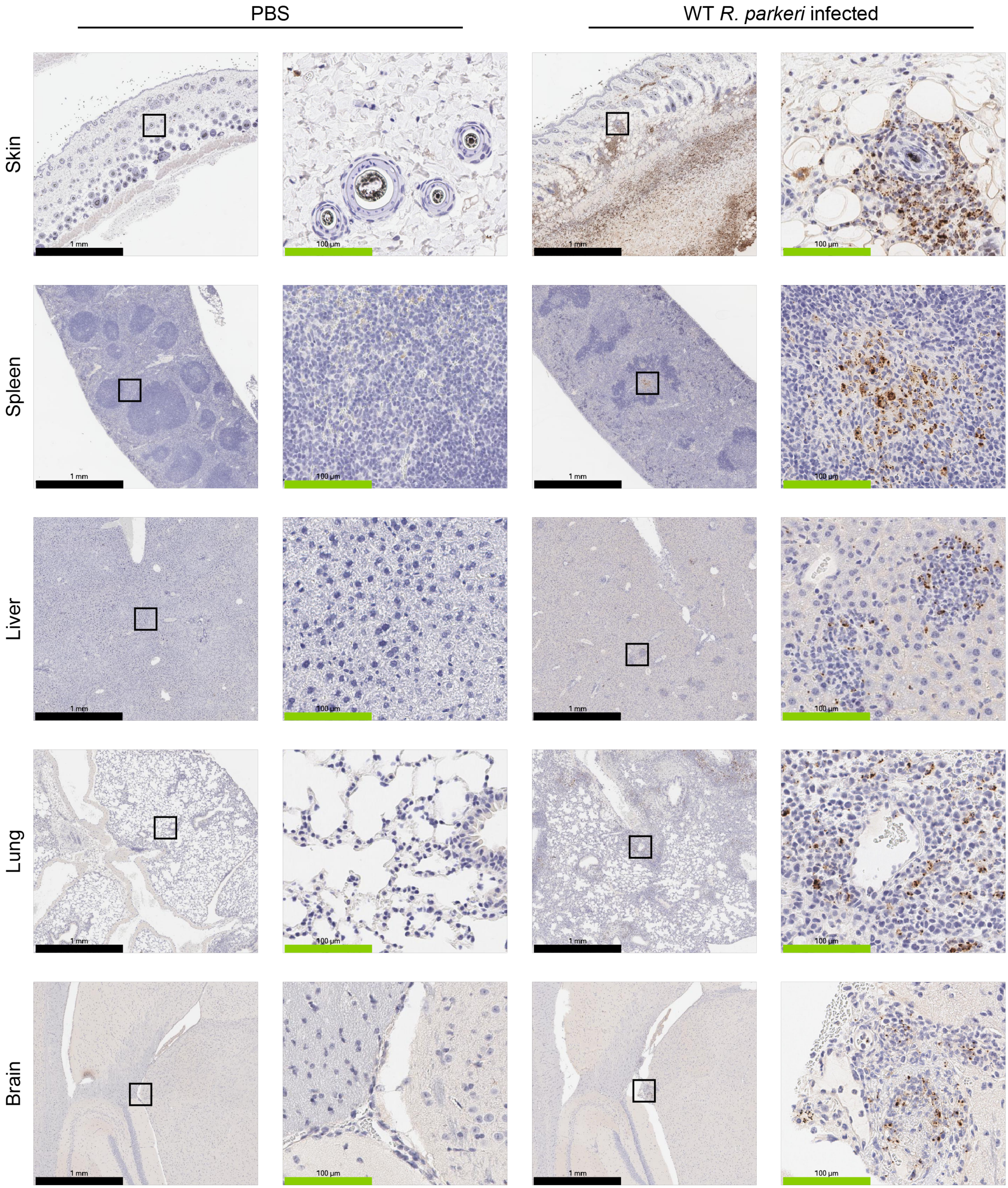


**Figure S3, related to Figure 7 – IHC staining of various organs.**

Representative images of anti-*Rickettsia* IHC staining of the indicated organs in mice either injected i.d. with PBS (control) or infected with $10^3$ pfu WT *R. parkeri*. Results were qualitatively similar for *rickA*::Tn-infected and *sca2*::Tn-infected mice. Black squares on the left were magnified on the right. Black scale bars represent 1 mm, green scale bars represent 100 μm.
